# The asymmetric transfers of visual perceptual learning determined by the stability of geometrical invariants

**DOI:** 10.1101/2024.01.02.573923

**Authors:** Yan Yang, Yan Zhuo, Zhentao Zuo, Tiangang Zhuo, Lin Chen

**Affiliations:** State Key Laboratory of Brain and Cognitive Science, Institute of Biophysics, Chinese Academy of Sciences, Beijing 100101, China; Hefei Comprehensive National Science Center, Institute of Artificial Intelligence, Hefei 230088, China; University of Chinese Academy of Sciences, Chinese Academy of Sciences, Beijing 100049, China

## Abstract

We quickly and accurately recognize the dynamic world by extracting invariances from highly variable scenes, a process can be continuously optimized through visual perceptual learning (VPL). While it is widely accepted that the visual system prioritizes the perception of more stable invariants, the influence of the structural stability of invariants on VPL remains largely unknown. In this study, we designed three geometrical invariants with varying levels of stability for VPL: projective (e.g., collinearity), affine (e.g., parallelism), and Euclidean (e.g., orientation) invariants, following the Klein’s Erlangen program. We found that learning to discriminate low-stability invariant transferred asymmetrically to those with higher stability, and that training on high-stability invariants enabled location transfer. To explore learning-associated plasticity in the visual hierarchy, we trained deep neural networks (DNNs) to model this learning procedure. We reproduced the asymmetric transfer between different invariants in DNN simulations and found that the distribution and time course of plasticity in DNNs suggested a neural mechanism similar to the reverse hierarchical theory (RHT), yet distinct in that invariant stability—not task difficulty or precision—emerged as the key determinant of learning and generalization. We propose that VPL for different invariants follows the Klein hierarchy of geometries, beginning with the extraction of high-stability invariants in higher-level visual areas, then recruiting lower-level areas for the further optimization needed to discriminate less stable invariants.

## Introduction

In order to adapt to a continually evolving and changing environment, the organism must identify information that remains consistent and stable in dynamic changes (***Gold and Stocker, 2017***). Through the process of visual perceptual learning (VPL), the visual system acquires an increased ability to extract meaningful and structured information from the environment to guide decisions and actions adaptively (***Adolph and Kretch, 2015***; ***Gibson and Pick, 2000***; ***Gibson, 1969***; ***Gold and Watanabe, 2010***). The “information” mentioned above can be conceptualized as invariance preserved under transformation in perception (***Gibson, 1979***).

It is widely accepted that different kinds of invariant properties hold distinct ecological significance and possess different levels of utility in perception (***Buccella, 2021***). A fairly large set of experimental findings across various paradigms have converged at the conclusion that the detectability and perceptual salience of an object’s attributes are systematically related to their structural stability under change—in a manner similar to Klein hierarchy of geometries; in particular, observers are more sensitive to geometrical properties with higher structural stability (***Chen, 2005***, ***1985***, ***1982***; ***Todd et al., 2014***, ***1998***). According to Klein’s Erlangen Program (***Klein, 1893***), a geometrical property is considered as an invariant preserved over a corresponding shape-changing transformation, the more general a transformation group, the more fundamental and stable the geometrical invariants over this transformation group. Within this hierarchy, structures that remain invariant under all projective transformations (projective geometry, e.g., maintaining collinearity) exhibit the highest stability. Less stable geometries arise by imposing additional constraints on the transformation group; for instance, affine geometry adds constraints to the projective group (e.g., maintaining parallelism), and Euclidean geometry introduces further constraints (e.g., maintaining length or orientation). Thus, the hierarchy of geometrical invariants is nested: projective transformations encompass both affine and Euclidean transformations, while affine transformations encompass Euclidean transformations (***Figure 1***). Despite this foundational understanding, it remains unclear whether the learning of invariants with different levels of stability follows a predictable pattern—a question central to our research. To address this, we must first grasp the characteristics of VPL.

**Figure 1.**
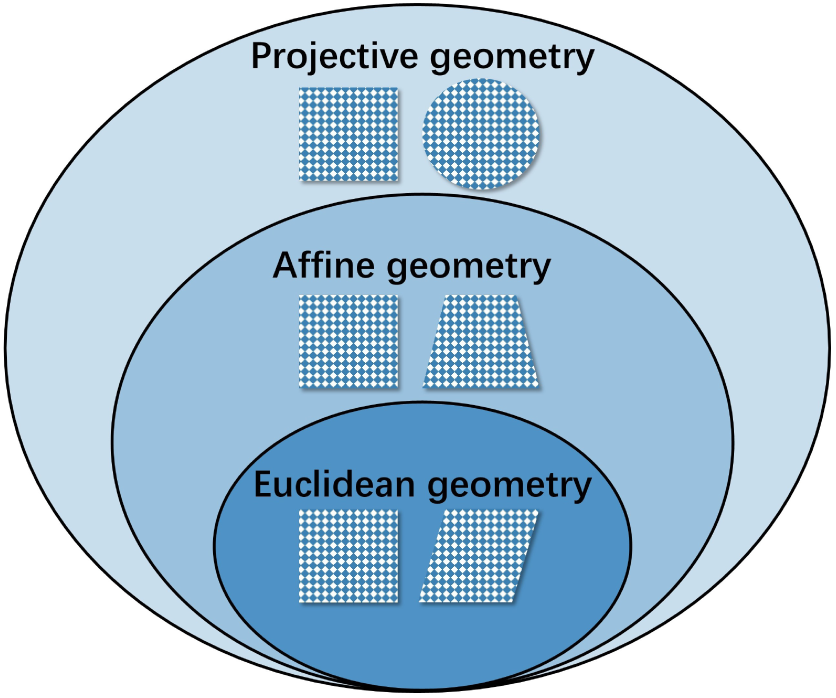
Nested relationship of the Klein hierarchy of geometries. Stratify geometrical invariants in ascending order of stability: Euclidean geometry, affine geometry, and projective geometry. The pairs of shapes in each circle differ in corresponding geometries.

According to Gibson’s differentiation view (***Gibson and Gibson, 1955***), perceptual learning is a process of “differentiating previously vague impressions” and whereby perceptual information becomes increasingly specific to the stimuli in the world. They proposed that learning through differentiation involves discovery and selectively processing the information most relevant to a task, including identifying “higher-order invariants” that govern some classification and filter out irrelevant information.

Traditionally, a hallmark of VPL has been its specificity for the basic attributes of the trained stimulus and task (***Crist et al., 1997***; ***Fiorentini and Berardi, 1981***; ***Hua et al., 2010***). Recent studies have challenged the specificity of learned improvements and demonstrated transfer effects between stimuli (***Liu and Weinshall, 2000***; ***Sowden et al., 2002***; ***Zhang et al., 2010***), location (***Hung and Seitz, 2014***) and substantially different tasks (***McGovern et al., 2012***; ***Szpiro and Carrasco, 2015***). To be of practical utility, the generalization of learning effects should be a research focus, and understanding the determinants of specificity and transfer remains one of the large outstanding questions in field. Ahissar and Hochstein uncovered the relationship between task difficulty and transfer effects (***Ahissar and Hochstein, 1997***), leading to the formulation of the reverse hierarchy theory (RHT) which suggests that VPL is a top-down process that originates from the top of the cortical hierarchy and gradually progresses downstream to recruit the most informative neurons to encode the stimulus (***Ahissar and Hochstein, 2004***). In this framework, the level at which learning occurs is related to the difficulty of the task: easier tasks are learned at higher-level visual areas and show more transfer, while harder tasks engage lower levels and are more specific. There is also evidence demonstrating greater specificity in finer precision tasks, suggesting that the precision of stimuli may be crucial in driving specificity (***Jeter et al., 2009***; ***Liu and Weinshall, 2000***; ***Hung and Seitz, 2014***). Another study (***Manenti et al., 2023***) introduced variability in a task-irrelevant feature during the training, finding that variability enables generalization of learning to new stimuli and locations, irrespective of the required precision of task. Furthermore, training with high-salience stimuli leads to plasticity in higher visual areas, thereby reducing specificity for spatial position and stimulus features (***Kourtzi et al., 2005***).

The research described in the present article concerns the very nature of form perception, trying to explore whether VPL of geometrical invariants with various stability also exhibit hierarchical relationships and what a priori rule defines the mode of learning and generalization. Our research focuses on the VPL of discriminations based on differences in geometrical invariants with different levels of structural stability: (1) projective property that is invariant over all projective transformations, such as whether a contour is straight or curved (collinearity); (2) affine property that is invariant under affine transformations, such as whether a pair of contours is parallel or nonparallel (parallelism); (3) Euclidean property that is only invariant under Euclidean transformation, such as the relative orientations of line segments (orientation). We conducted three psychophysics experiments assessing how the structural stability of geometrical properties affect the learning effect, meanwhile investigating the transfer effect between different geometrical invariants or between different locations within each invariant. We modeled the learning processes using a deep neural network (DNN) that recapitulates several known VPL phenomena and thus provides a promising testbed to study learning-related plasticity in the primate visual hierarchy (***Wenliang and Seitz, 2018***). The learning-induced changes in the DNN model yield important predictions to the underlying neural substrate. We then interpret the results based on the Klein hierarchy of geometries and RHT.

## Results

### Asymmetric transfer effect: The learning effect consistently transferred from low-stability to high-stability invariants

The paradigm of “configural superiority effects” with reaction time measures Forty-four right-handed healthy subjects participated in Experiment 1. We randomly assigned subjects into three groups trained with one invariant discrimination task: the collinearity (colli.) training group (n = 15), the parallelism (para.) training group (n = 15) and the orientation (ori.) training group (n = 14). The paradigm of “configural superiority effects” (***Chen, 2005***; ***Pomerantz et al., 1977***) (faster and more accurate detection/discrimination of a composite display than of any of its constituent parts) was adapted to measure the short-term perceptual learning effects of different levels of invariants. As illustrated in ***Figure 2A***, subjects performed the odd-quadrant discrimination task in which they were asked to report which quadrant differs from the other three as fast as possible on the premise of accuracy. During the test phases before and after training (Pre-test and Post-test, ***Figure 2B***), subjects performed the three invariant discrimination tasks (colli., para., ori.) at three blocks respectively, the response times (RTs) were recorded to measure the learning effects. To distinguish VPL from programmed learning due to motor learning, a color discrimination task, served as baseline training, were performed before the main experiment (***Figure 2B***).

**Figure 2.**
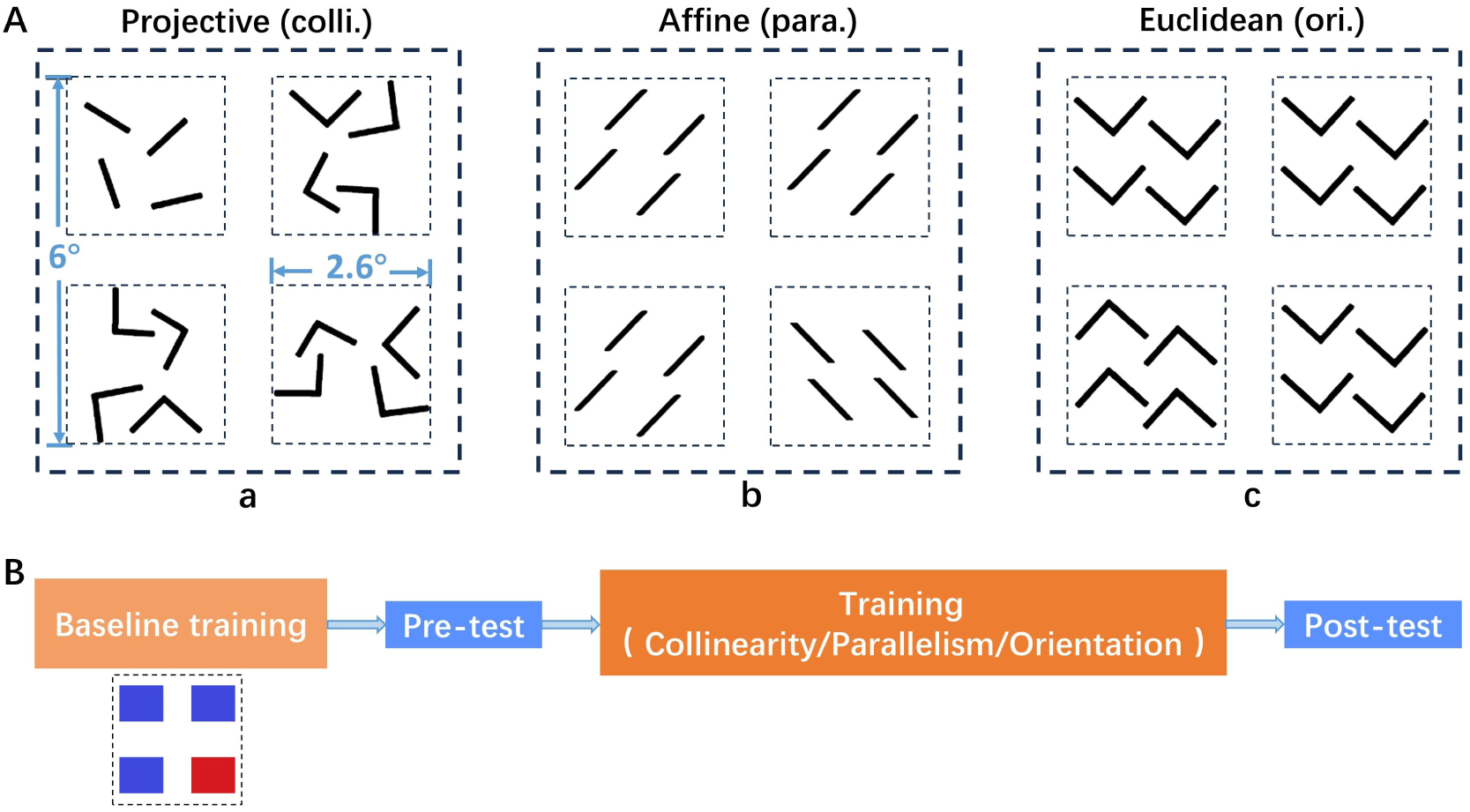
(A) Examples of stimulus arrays designed to measure the perceptual learning effects of the geometries at different levels of structural stability in an odd quadrant task. They represent discriminations based on (a) a difference in collinearity, a kind of projective property, (b) a difference in parallelism, a kind of affine property, and (c) a difference in angle orientation, a kind of Euclidean property. (B) Procedure of Experiment 1. **Figure 2—figure supplement 1**. The configural superiority effect.

First of all, to investigate if there was any speed-accuracy trade-off, one-way repeated measures analysis of variance (ANOVA) with task as within-subject factors was conducted on the accuracies collected in the Pre-test phase. A significant main effect of the task was found (F(2, 129) = 7.977, p = 0.0005, 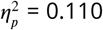). We performed further post-hoc analysis carrying out paired t-test with FDR correction to examine the differences of accuracies between each pair of tasks. As a result, the accuracies of the collinearity task were significantly higher than that in the parallelism task (t(43) = 5.443, p < 0.0001, Hedge’s g = 0.917), and that in the orientation task (t(43) = 4.351, p = 0.0001, Hedge’s g = 0.574). There was no difference in accuracy between the parallelism and orientation task (t(43) = 1.535, p = 0.132, Hedge’s g = 0.214). Then the same analysis was applied to the RTs in the three tasks prior to training. A significant main effect of the task was found in ANOVA (F(2, 129) = 59.557, p < 0.0001, 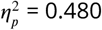). As shown from the post-hoc test, the RTs of collinearity task were significantly faster than that in the parallelism task (t(43) = 13.374, p < 0.0001, Hedge’s g = 1.945), and that in the orientation task (t(43) = 13.333, p < 0.0001, Hedge’s g = 2.295). The RT of the parallelism task was faster than that of orientation task (t(43) = 4.179, p < 0.0001, Hedge’s g = 0.416). Taken together, the collinearity task has the highest accuracy as well as the faster RT among the three tasks, showing no speed-accuracy trade-off in Experiment 1. What’s more, the results before training were in line with the prediction from the Klein hierarchy of geometries, which suggest that the more stable invariants possessed higher detectability, resulting in better task performance. The statistical results of the accuracies in Post-test didn’t differ from that in Pre-test (***Figure 3—figure Supplement 1***). In the following analysis, only correct trials were used.

**Figure 3.**
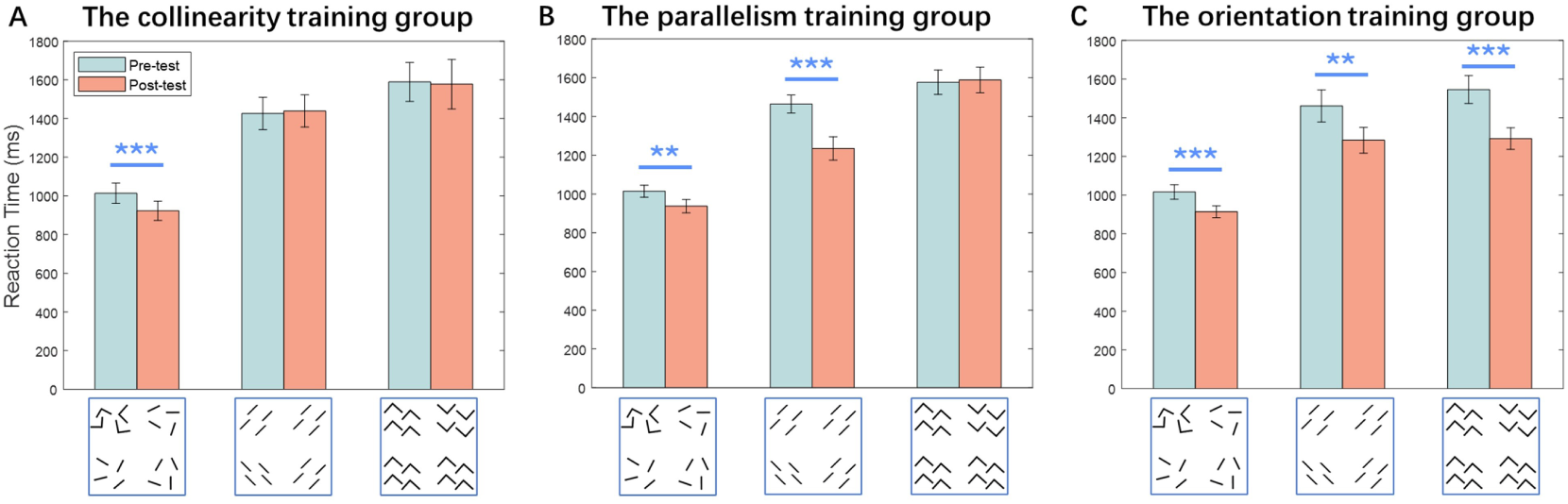
Results of Experiment 1, RTs of each discrimination tasks measured at Pre-test and Post-test were compared by one-tailed, paired t-test. (A) Results from the group trained on the collinearity task (n=15). Performances of the collinearity task were improved after training (p = 0.0008). (B) Results from the group trained on the parallelism task (n=15). Performances of the collinearity (p = 0.008) and parallelism (p < 0.0001) task were improved after training. (C) Results from the group trained on the orientation task (n=14). Performances of the collinearity (p = 0.0007), parallelism (p = 0.002) and orientation task (p = 0.0002) were improved after training. (***p < 0.001, **p < 0.01, *p < 0.05). Error bars denote 1 SEM across subjects. **Figure 3—figure supplement 1**. Accuracies for the three discrimination tasks measured at Pre-test and Post-test. **Figure 3—figure supplement 2**. The learning indexes of the three geometrical invariants in Experiment 1. **Figure 3—source data 1**. RTs and accuracies at Pre-test and Post-test, and learning indexes in the course of training for each participant.

The second analysis conducted was to assess whether there were learning effects of the trained tasks and transfer effects to the untrained tasks. This was assessed by examining the RTs of Pre-test and Post-test for each geometrical property discrimination task. One-tailed, paired t-test was performed to do this. For the collinearity training group, significant learning effect was found (t(14) = 3.911, p = 0.0008, Cohen’s d = 0.457), but there was no transfer to the other two untrained task (***Figure 3A***). The parallelism training group show significant learning effect (t(14) = 5.169, p < 0.0001, Cohen’s d = 1.095), and also show substantial improvement in the collinearity task (t(14) = 2.753, p = 0.008, Cohen’s d = 0.609) (***Figure 3B***). Moreover, performances of all three task were improved after training on orientation discrimination task: the collinearity task (t(13) = 4.033, p = 0.0007, Cohen’s d = 0.800), the parallelism task (t(13) = 3.482, p = 0.002, Cohen’s d = 0.631), and the orientation task (t(13) = 4.693, p = 0.0002, Cohen’s d = 1.048) (***Figure 3C***). This particular pattern of transfer is interesting given the hierarchical relationship of the three different stimulus configurations. For instance, the performance improvement obtained on the parallelism task transferred to the collinearity task which is more stable, whereas not transferred to the orientation task which is less stable. Similarity, learned improvement in orientation discrimination transferred to more stable tasks, with RTs of both collinearity and parallelism tasks showing significant improvements. However, training on collinearity discrimination which is the most stable among the three tasks exhibited task specificity. These findings indicate that perceptual improvements derived from training on a relatively low-stability form invariant can transfer to those invariants with higher stability, but not vice versa.

Finally, we assessed whether learning effect differ in different form invariants. To this end, we computed the “learning index” (LI) (***Petrov et al., 2011***), which quantifies learning relative to the baseline performance. ANOVA analysis did not find a significant difference in the learning indices among the three tasks (F(2, 41) = 2.246, p = 0.119, 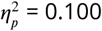) (***Figure 3—figure Supplement 2***).

What needs to be cautious is that, Experiment 1 with RT measures has limitations that make it difficult to truly compare the time required for processing different invariants. Specifically, our interest lies in understanding how learning affects the process of extracting geometrical invariants. However, in order to make a response, participants also need to locate the differing quadrant. The strength of the grouping effect of the shapes among the four quadrants can affect the speed of the localization process (***Orsten-Hooge et al., 2011***). Additionally, the strength of the grouping effect may vary under different conditions, leading to differences in reaction times that may reflect differences in the extraction time of geometrical invariants as well as the strength of the group effect among the quadrants.

The paradigm of “configural superiority effects” with reaction time measures To overcome the shortcomings of the RT measures, VPL is indexed by the improvements in thresholds of discrimination tasks after training in Experiment 2. We employed the adaptive staircase procedure QUEST (***Watson and Pelli, 1983***) to assess the thresholds. QUEST is a kind of Bayesian adaptive methods which typically produces estimates of psychometric function parameters and converges to a threshold more quickly than conventional staircase procedures. Forty-five healthy subjects participated in Experiment 2, and they were randomly assigned into three groups: the collinearity training group (n = 15), the parallelism training group (n = 15) and the orientation training group (n = 15). On each trial, subjects are required to determine whether a “target” is present among two simultaneously displayed stimuli (first-order choice, detection task), and if it is present, further specify its position (second-order choice, two-alternative forced choice task) (***Figure 4***, ***Figure 5*** and ***Figure 4—figure Supplement 1***).The procedure of Experiment 2 is similar to that of Experiment 1 except for no involvement of the baseline training. For each block, a QUEST staircase was used to adaptively adjust the angle separation (*θ*) of discrimination task within all trials but the catch trials, and provided an estimate of each subject’s 50% correct discrimination threshold.

**Figure 4.**
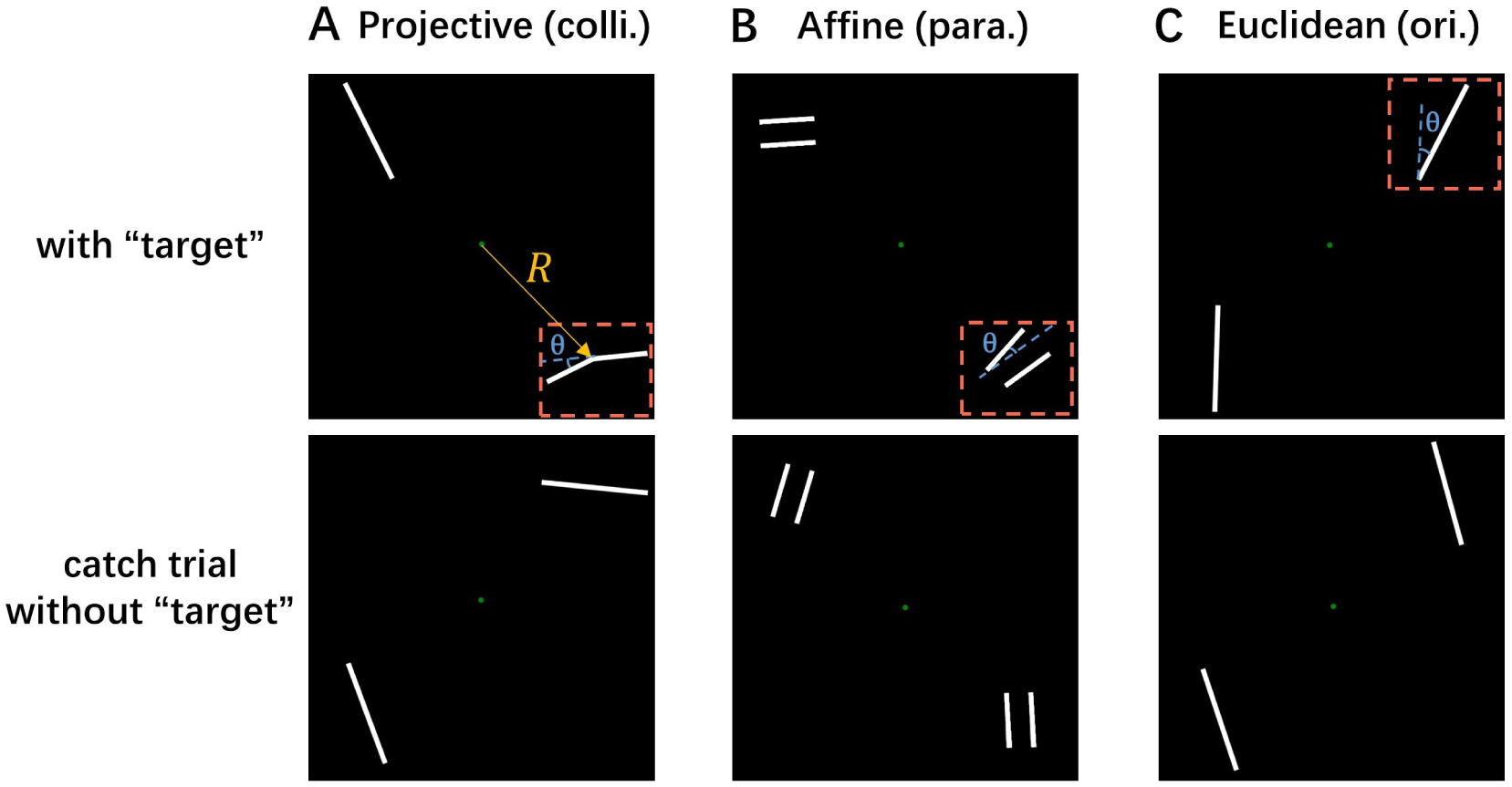
Examples of the layout of a stimulus frame with two stimuli presented in diagonally opposite quadrants. The top row demonstrates trials with a “target” (surrounded by orange dashed box), and the bottom row demonstrates the catch trials without “target”. The blue dashed lines represent the “base” orientation for each stimulus, and *θ* is the angle separation of the discrimination task. (A) Stimulus examples of the collinearity (colli.) task, the upper example shows a “target” (a line with one bend along its length) located at the lower right quadrant. Two straight lines were presented on a catch trial. (B) Stimulus examples of the parallelism (para.) task, the upper example shows a “target” (a pair of unparallel lines) located at the lower right quadrant. Two pairs of parallel lines were presented on a catch trial. (C) Stimulus examples of the orientation (ori.) task, the upper example shows a “target” (the more clockwise line) located at the upper right quadrant. Two lines with identical orientation were presented on a catch trial. **Figure 4—figure supplement 1**. Examples of stimuli in Experiment 2.

**Figure 5.**
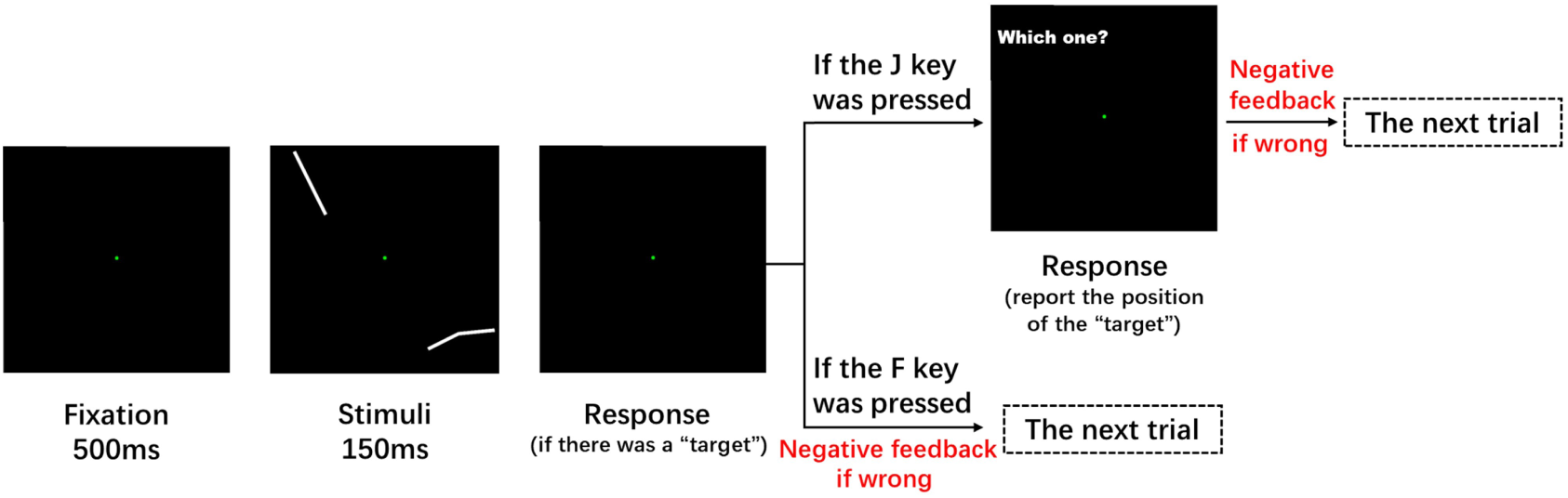
Schematic descriptions of a trial in Experiment 2. A trial included three intervals: a 500-ms pre-stimulus epoch, a 150-ms stimulus epoch, and the response epoch. In the stimulus epoch, two stimuli were presented at two diagonally opposite quadrants. Subjects maintained fixation in a green fixation point until the stimulus disappear, and indicated if there was a “target” by pressing the corresponding key (“J” for Yes, “F” for No). They should further report the position of the “target” if they pressed the J key. Negative feedback tone was presented at the end of a trial if incorrect response was given.

One-way repeated measures ANOVA and post-hoc t-test were conducted on the thresholds collected from all three training groups prior to training. A significant main effect of the task was found in ANOVA analysis (F(2, 132) = 13.598, p < 0.0001, 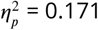). As revealed by post-hoc tests, the initial performances of the three tasks were consistent with the relative stability of the invariant they involved, the discrimination threshold of the collinearity task was significantly lower than that of the parallelism task (t(44) = 3.247, p = 0.002, Hedge’s g = 0.595), and that of the orientation task (t(44) = 4.225, p = 0.0001, Hedge’s g = 0.871). The threshold of the parallelism task was lower than that of the orientation task (t(44) = 3.267, p = 0.002, Hedge’s g = 0.676). Moreover, the accuracies of the three tasks in Pre-test were also submitted to a one-way repeated measures ANOVA and no significant effect was found (F(2, 132) = 0.046, p = 0.955, 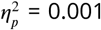), suggesting no difference in difficulty among the three tasks before training.

One-tailed, paired t-test was performed to compare the threshold in Pre-test and Post-test. After orientation discrimination training, the orientation discrimination threshold at Post-test was significantly lower than that at Pre-test (t(14) = 2.443, p = 0.014, Cohen’s d = 0.898), and the same applied to the collinearity discrimination task, t(14) = 2.740, p = 0.008, Cohen’s d = 0.752, and the parallelism discrimination task, t(14) = 1.949, p = 0.036, Cohen’s d = 0.654 (***Figure 6C***). After parallelism discrimination training, significant improvements were found in the collinearity discrimination task (t(14) = 2.775, p = 0.007, Cohen’s d = 1.013) and the parallelism discrimination task (t(14) = 3.259, p = 0.003, Cohen’s d = 1.192) (***Figure 6B***). And collinearity discrimination training only produced an improvement on its own performance (t(14) = 2.128, p = 0.026, Cohen’s d = 0.759, ***Figure 6A***). In summary, the pattern of generalization is identical to what was found in Experiment 1, where training of low-stability invariants optimized the perception of high-stability invariants but not vice versa. Just the same as in Experiment 1, no differences were found between the LIs of the three tasks in Experiment 2 (F(2, 42) = 2.905, p = 0.066, 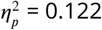, ***Figure 6—figure Supplement 1***).

**Figure 6.**
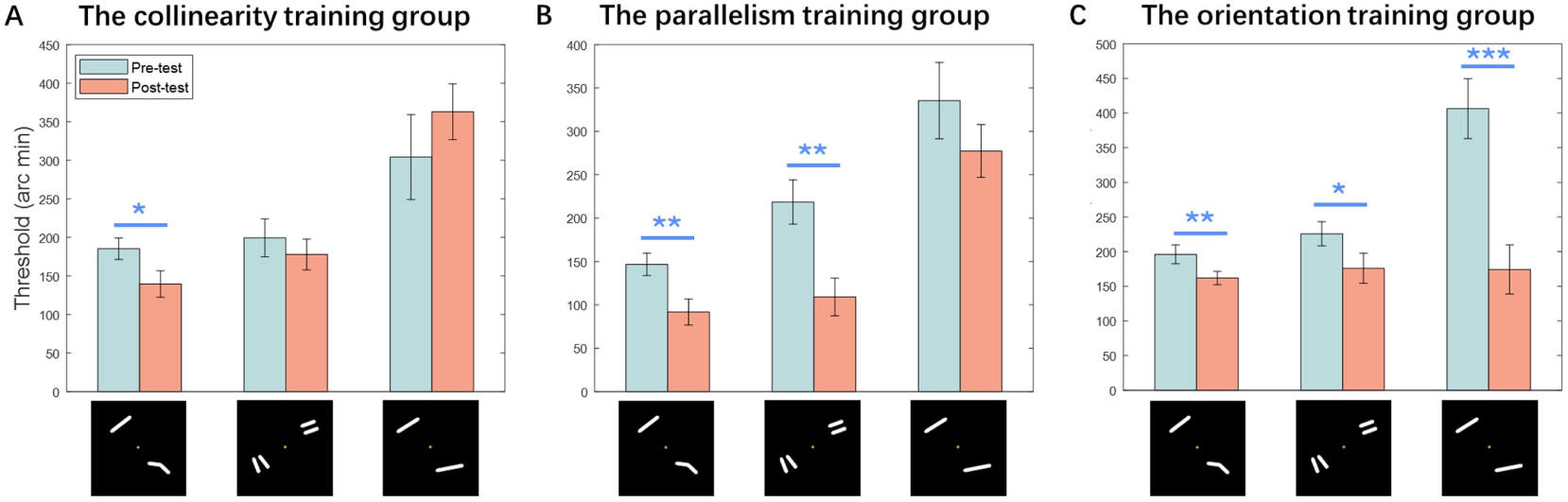
Results of Experiment 2, Thresholds of each discrimination task measured at Pre-test and Post-test were compared by one-tailed, paired t-test. (A) Results from the group trained on the collinearity task (n = 15). Performances of the collinearity task were improved after training (p = 0.026). (B) Results from the group trained on the parallelism task (n = 15). Performances of the collinearity (p = 0.007) and parallelism (p = 0.003) task were improved after training. (C) Results from the group trained on the orientation task (n = 15). Performances of the collinearity (p = 0.008), parallelism (p = 0.036) and orientation task (p < 0.0001) were improved after training. (***p < 0.001, **p < 0.01, *p < 0.05). Error bars denote 1 SEM across subjects. **Figure 6—figure supplement 1**. The learning indexes of the three geometrical invariants in Experiment 2. **Figure 6—figure supplement 2**. Long-term learning curves across the three invariant discrimination tasks. **Figure 6—source data 1**. Thresholds and accuracies at Pre-test and Post-test, and learning indexes in the course of training for each participant. **Figure 6—source data 2**. Thresholds in the course of long-term learning for each participant.

The employment of short-term perceptual learning paradigms has made it difficult to accurately track the temporal learning processes (***Yang et al., 2022***).It is also possible that the lack of observed differences in learning effects between tasks could be due to insufficient learning in some tasks, considering the possibility that different training tasks may involve different short-term and longterm learning processes (***Aberg et al., 2009***; ***Mascetti et al., 2013***). Given the considerations above, we recruited three additional groups of subjects (n=7 for each group) for long-term training consisting of five consecutive daily sessions. We computed bootstrapped estimates of the learning curve for each task because of the small sample size (***Figure 6—figure Supplement 2***). Power parameter (the ‘*b*’ parameter for *y* = *ax*^*b*^) estimates for the groups trained on colli., para. and ori. Were −0.076 (95% CI [−0.159, 0.008]), −0.136 (95% CI [−0.229, −0.034]), and −0.223 (95% CI [−0.372, −0.156]), respectively, suggesting that learning rates (equivalent to magnitudes of learning) are higher for less stable invariants.

Previous studies claimed that transfer of VPL is controlled by the difficulty or precision of the training task (***Ahissar and Hochstein, 1997***; ***Jeter et al., 2009***; ***Wenliang and Seitz, 2018***). In this experiment, task difficulty is related to the accuracy, and task precision which is related to the angle separation can be indexed by the threshold. As stated above, prior to training, there was not significant difference among the difficulties (accuracies) of the three tasks, and tasks with higher stability had lower threshold values, resulting in higher precision during training. We showed that the relative stability of invariants determined the transfer effects between tasks even when task difficulty was held constant between tasks, and the particular transfer pattern found in our study (learning from tasks with lower stability and lower precision transferred to the tasks with higher stability and precision) is contrary to the precision-dependent explanation for the generalization of VPL which proposed that training on higher precision can improve performance on lower precision tasks but the reverse is not true (***Jeter et al., 2009***; ***Wenliang and Seitz, 2018***).

### Location generalization within each invariant

Having revealed the asymmetric transfer of learning between geometrical invariants with varying levels of stability, we then consider the extent to which learning transfers across locations within each invariant condition in Experiment 3. Experiment 2 was adapted for testing this within-invariant transfer effect. Thirty-eight healthy subjects participated in Experiment 3, and they were randomly assigned into three groups: the collinearity training group (n = 13), the parallelism training group (n = 12) and the orientation training group (n = 13). Each group of subjects performed their assigned task on fixed location during the training phase, and then were tested on the same task with stimuli presented at new location (the transfer test). Specifically, if the phase of training used the upper-left/lower-right diagonal, then the transfer test used the upper-right/lower-left diagonal, and vice versa. The presentation diagonal was randomly assigned to subjects for initial training and switched to the opposite diagonal for the transfer test.

We assessed learning (first vs. last trained block) and transfer effects (first trained block vs. transfer test) using one-tailed, paired t-test. All three groups demonstrated statistically significant increases in discrimination performance after training (colli.: t(12) = 7.821, p < 0.0001, Cohen’s d = 2.032; para.: t(11) = 7.688, p < 0.0001, Cohen’s d = 1.893; ori.: t(12) = 6.153, p < 0.0001, Cohen’s d = 1.931), along with significant transfer effects to untrained location (colli.: t(11) = 5.324, p = 0.0001, Cohen’s d = 1.372; para.: t(11) = 3.704, p = 0.002, Cohen’s d = 0.809; ori.: t(12) = 3.641, p = 0.002, Cohen’s d = 0.948) (***Figure 7A***).

**Figure 7.**
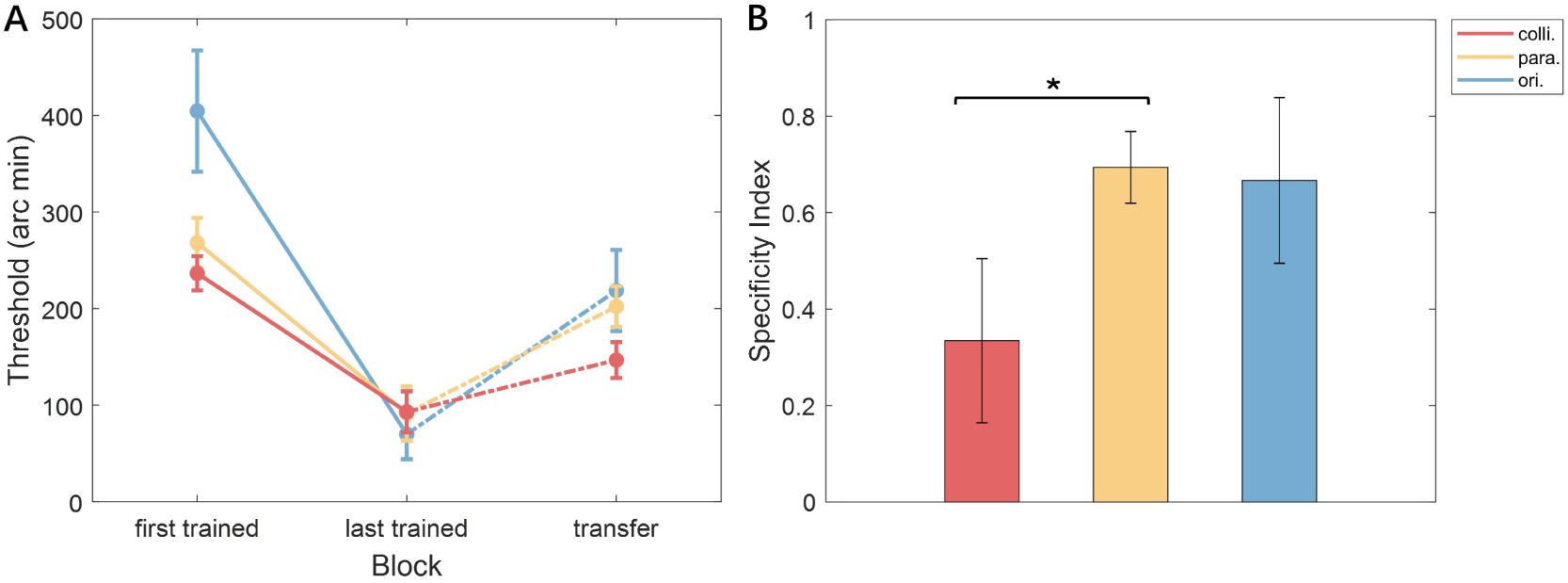
Results of Experiment 3. (A) Thresholds of each discrimination task measured at the first trained block, the last trained block, and the transfer test. (B) The specificity indexes of the three geometrical invariants. Two-tailed, two-sample t-test were conducted to compare the SIs between each pair of the three tasks. Significant difference was only found between collinearity and parallelism task (p = 0.020). Error bars denote 1 SEM across subjects. **Figure 7—source data 1**. Thresholds at the first trained block, the last trained block and the transfer test for each participant.

To further assess the location transfer effect of VPL for the three groups, we computed the “specificity index” (SI) (***Ahissar and Hochstein, 1997***), which estimates the portion of the initial learning that is transferred to untrained conditions (***Figure 7B***). Positive SI values indicate that learning is specific, whereas SI values smaller than or equal to 0 indicate generalization. We used two-tailed, one-sample t-test for comparisons against 0 or 1. For the collinearity training group, the SIs were not significantly above 0 (t(12) = 2.042, p = 0.064, Cohen’s d = 0.566) but significantly below 1 (t(12) = −5.069, p = 0.0003, Cohen’s d = 1.406), demonstrating generalization to new location. Partial location transfer was found in the parallelism and orientation task, for which the SIs were significantly above 0 (para.: t(11) = 9.324, p < 0.0001, Cohen’s d = 2.692; ori.: t(12) = 4.526, p = 0.001, Cohen’s d = 1.180) and significantly below 1 (para.: t(11) = −4.114, p = 0.002, Cohen’s d = 1.188; ori.: t(12) = −2.789, p = 0.016, Cohen’s d = 0.774). Furthermore, significant difference in SIs was observed only between the collinearity and parallelism training groups (two-tailed, two-sample t-test: t(18.112) = −2.556, p = 0.020, Cohen’s d = 0.998), indicating that the collinearity task exhibits greater location generalization compared to the parallelism task.

The location transfer observed in the orientation task is somewhat surprising, given that VPL of orientation discrimination is typically reported to be highly specific to the trained retinal location (***Fiorentini and Berardi, 1980***; ***Schoups et al., 1995***; ***Shiu and Pashler, 1992***). Although previous studies have observed location transfer in orientation discrimination with low precision (large angle separations), our study’s angle separation thresholds were closer to those associated with location specificity (***Jeter et al., 2009***; ***Manenti et al., 2023***). We therefore speculate that the observed location transfer in the orientation task may result from our unique experimental design: in the orientation task, subjects needed to compare pairs of lines presented in different quadrants to identify the “target” (***Figure 4***), whereas in the collinearity and parallelism tasks, subjects could make decisions based on stimuli within a single quadrant. This requirement for spatial integration in the orientation task likely enhanced its location generalization.

Overall, the greater location transfer observed in the collinearity task compared to the parallelism task, along with the location specificity in orientation discrimination reported in previous VPL studies, suggests that training with more stable invariants reduces spatial specificity. This finding also implies that plasticity associated with more stable invariants likely occurs in higher-order visual cortex areas, where neurons have larger receptive fields. As in Experiment 2, neither task difficulty nor stimulus precision can account for the location generalization observed in our study.

### Deep neural network simulations of learning and transfer effects

#### Behavioral results

In Experiment 4, we repeated Experiment 2 using a DNN to model VPL (***Wenliang and Seitz, 2018***), aiming to determine whether training the DNN with invariants yields comparable behavioral improvements to those observed in humans and to gain insight into the underlying neural mechanisms. This network, which is derived from the general AlexNet architecture, fulfilled predictions of existing theories (e.g. the RHT) regarding specificity and plasticity and reproduced findings of tuning changes in neurons of the primate visual areas (***Manenti et al., 2023***; ***Wenliang and Seitz, 2018***). Three networks were trained on the three tasks (colli., para., ori.) respectively, repeated in 12 conditions with varying stimulus parameters.

The performance trajectories of the networks trained with collinearity, parallelism, and orientation discrimination were averaged across the 12 stimulus conditions, and are shown in ***Figure 8A*** respectively. Each network was also tested on the two untrained tasks in the same stimulus condition during the training phase, and the transfer accuracies are presented in ***Figure 8A*** as well. As shown from the final accuracies (the numbers located at the end of each curve), the networks trained on ori. showed greatest transfer effects to the untrained tasks, and the networks trained on colli. showed worst transfer effects.

**Figure 8.**
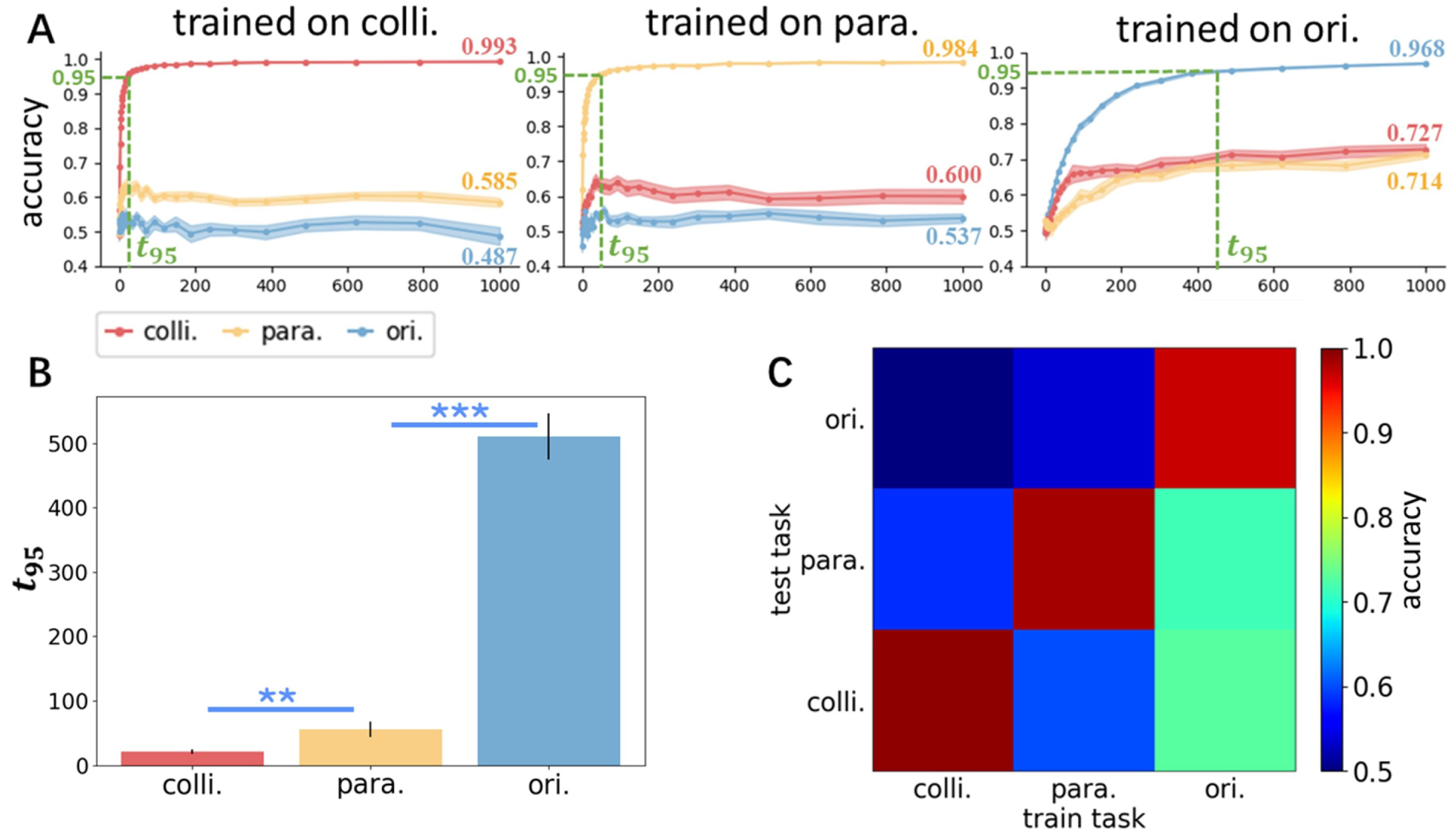
Performance of the model when trained under different discrimination tasks. (A) Accuracy trajectories against training iterations from the models trained on collinearity (left), parallelism (middle), and orientation task (right), with the error bar representing 1 SEM. *t*_95_ is the iteration where the fully plastic network reached 95% accuracy, depicted by green dashed lines. The numbers located at the end of each curve are the final accuracies of the last iteration. (B) The learning speed which was indexed by *t*_95_ of the three tasks. The learning speed of the collinearity task was faster than the parallelism (p = 0.018) and orientation task (p < 0.0001). The learning speed of the parallelism task was faster than the orientation task (p < 0.0001). Statistical significance was calculated by paired t-test with FDR correction. (***p < 0.001, **p < 0.01, *p < 0.05). Error bars denote 1 SEM across subjects. (C) Final mean accuracies when the network was trained and tested on all combinations of tasks **Figure 8—figure supplement 1**. Stimulus examples in Experiment 3.

We then investigate the speeds of learning of the three tasks, we calculated *t*_95_ (***Wenliang and Seitz, 2018***), the iteration where the fully plastic network reached 95% accuracy, for each task. We found a significant main effect of the training task on *t*_95_ using the one-way repeated measures ANOVA (F(2, 33) = 144.636, p < 0.0001, 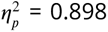). Further post-hoc analysis with paired t-test showed that the performance of collinearity discrimination started to asymptote earlier than the parallelism (t(11) = 2.567, p = 0.018, Cohen’s d = 1.012) and the orientation task (t(11) = 13.172, p < 0.0001, Cohen’s d = 5.192). The orientation discrimination had slowest speed of learning, it reached saturation slower than the parallelism task (t(11) = 11.626, p < 0.0001, Cohen’s d = 4.583) (***Figure 8B***).

***Figure 8C*** shows the final learning and transfer performance on all combinations of training and test task. A linear regression on the final accuracies showed a significant positive main effect of the stability of test task on the performance (*β* = 0.055, t(105) = 2.467, p = 0.015, *R*^2^ = 0.043), shown as increasing color gradient from top to bottom. We also found that the stability of training task had a significant negative effect (*β* = −0.057, t(105) = −2.591, p = 0.011, *R*^2^ = 0.048), shown as decreasing color gradient from right to left. Overall, consistent with the results in the two psychophysical experiments, these results suggested that transfer is more pronounced from less stable geometrical invariants to more stable invariants than vice versa, shown as higher accuracy on lower-right quadrants compared with top-left quadrants.

#### Distribution of learning across layers

To demonstrate the distribution of learning over different levels of hierarchy, we next examined the time course of learning across the layers, the weight changes for each layer were shown in ***Figure 9A***. Overall, training on lower-stability geometrical invariants produced greater overall changes. Due to this mismatch of weight initialization, we focus on layers 1–5 with weights initialized from the pre-trained AlexNet.

**Figure 9.**
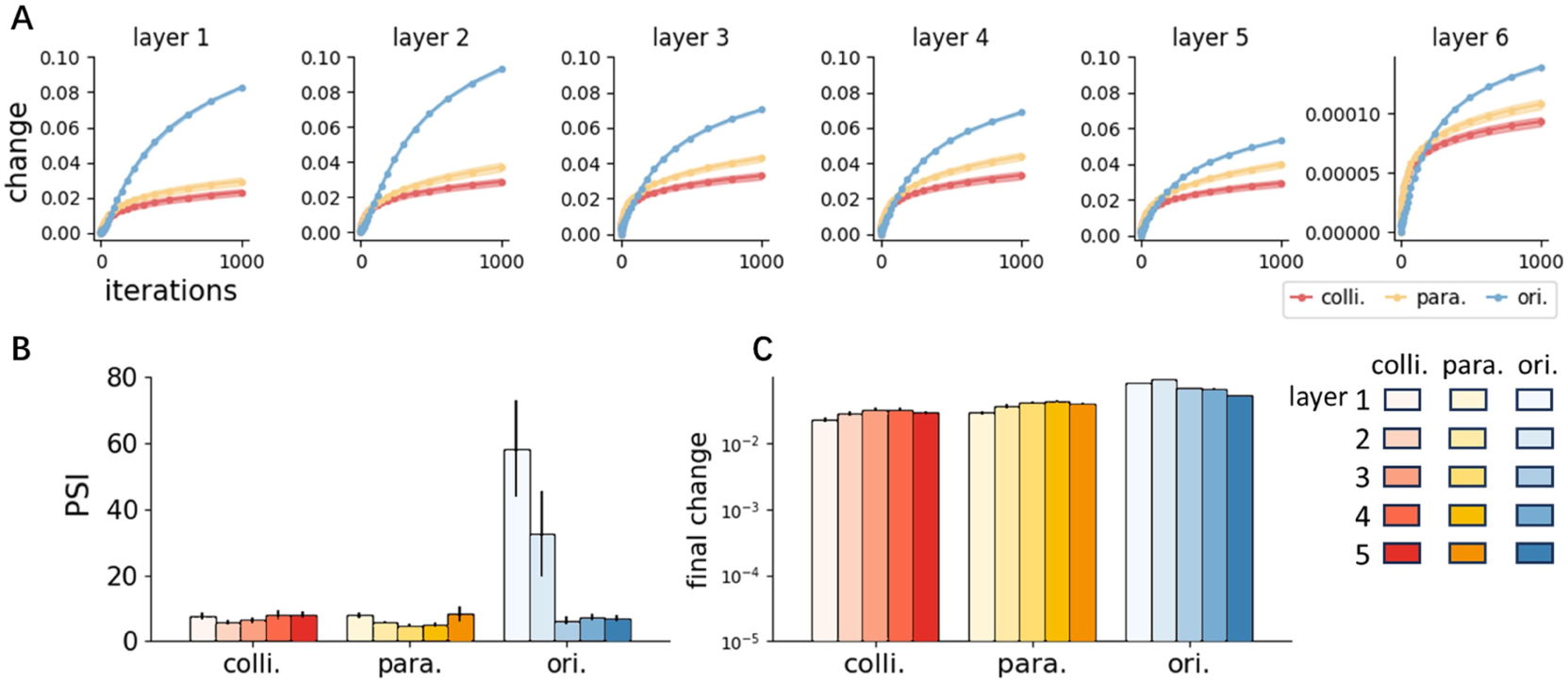
Layer change under different training tasks. (A) Layer change trajectories during learning. (B) Iteration at which the rate of change peaked (PSI) in layers 1-5. (C) Final layer change in layers 1-5. The error bar representing 1 SEM.

To characterize learning across layers, we studied when and how much each layer changed during training. First, to quantify when significant learning happened in each layer, we estimated the iteration at which the gradient of a trajectory reached its peak (peak speed iteration, PSI; shown in ***Figure 9B***). As the result of a linear regression analysis, in layers 1–5, we observed significant negative main effects of the stability of training task (*β* = −27.315, t(176) = −6.623, p < 0.0001, *R*^2^ = 0.072), layer number (*β* = −17.327, t(176) = −6.450, p < 0.0001, *R*^2^ = 0.065) and a positive interaction of the two on PSI (*β* = 6.579, t(176) = 5.290, p < 0.0001, *R*^2^ = 0.115), suggesting that layer change started to asymptote later for lower layers and less stable invariants. For individual tasks, a linear regression analysis showed a significant negative effect of layer number on PSI only in the least stable task, that is the orientation discrimination task (*β* = −12.833, t(58) = −4.470, p < 0.0001, *R*^2^ = 0.243). Therefore, for the discrimination of the least stable invariants, the order of change across layers is consistent with the RHT prediction that higher visual areas change before lower ones (***Ahissar and Hochstein, 2004***, ***1997***).

The final layer changes at the end of training for the networks trained on the three tasks are shown in ***Figure 9C*** respectively. A linear regression analysis on the final changes in layers 1–5 revealed significant negative main effects of the stability of training task (*β* = −0.037, t(176) = −16.409, p < 0.0001, *R*^2^ = 0.689), layer number (*β* = −0.011, t(176) = −7.655, p < 0.001, *R*^2^ = 0.0001) and a positive interaction of the two (*β* = 0.005, t(176) = 7.329, p < 0.0001, *R*^2^ = 0.075). However, for individual tasks, a significant negative linear effect of layer number was only found in the orientation discrimination task (*β* = −0.008, t(58) = −12.763, p < 0.0001, *R*^2^ = 0.733). On the contrary, significant positive effect of layer number was found in the parallelism (*β* = 0.003, t(58) = 3.701, p = 0.0005, *R*^2^ = 0.177) and collinearity task (*β* = 0.002, t(58) = 2.538, p = 0.014, *R*^2^ = 0.084). We then calculated the centroid, which is equal to the weighted average of the layer numbers using the corresponding mean final change as the weight. The centroids for the networks trained on colli., para. and ori. are 3.11, 3.14 and 2.77, respectively. These centroids together with the results from linear regression collectively indicate that the least stable task induces more change lower in the hierarchy whereas the two more stable tasks induce change higher in the hierarchy.

Taken together, these findings demonstrate that training with lower-stability invariants delayed the onset of asymptotic learning while resulting in greater overall changes across DNN layers by the end of training. For tasks involving more stable invariants, such as collinearity and parallelism, learning-induced changes occurred predominantly in higher layers. In contrast, the least stable tasks (orientation) primarily involved changes in lower layers.

The distribution of learning across DNN layers suggests that the behavioral VPL effects observed in both humans and DNNs can be understood within the framework of RHT. Specifically, VPL of more stable invariants likely occurs in higher-order visual areas, facilitating greater location generalization, as observed in Experiment 3. In contrast, VPL of less stable invariants requires further optimization where lower-order areas are involved. This results in a greater amount of learning for both humans (***Figure 6—figure Supplement 2***) and DNNs (greater final layer change) while producing slower but more substantial changes in lower layers (***Figure 9B, C***). More importantly, RHT also predicts a unidirectional transfer, where higher levels of the cortical hierarchy benefit from modifications of neural responses at lower levels (***Ahissar and Hochstein, 2004***, ***1997***; ***McGovern et al., 2012***). This is consistent with the asymmetric transfer from low- to high-stability invariants observed in our study.

Wenliang and Seitz proposed that high-precision training transfers more broadly to untrained and coarse discriminations than low-precision training (***Wenliang and Seitz, 2018***). However, in each stimulus condition of Experiment 4, the angle separations (precisions) in the collinearity and parallelism task were always the same, and were half of the angle separations in the orientation task. So, the pattern of transfer and layer change cannot be explained based on the relative precision of the training and test tasks that was suggested by Wenliang and Seitz. Rather, the relative stability of invariants involved in the tasks provides consistent and reasonable explanation for the asymmetric transfers found in our study.

## Discussion

In this study, we investigated perceptual learning of the three geometrical invariants in the Klein hierarchy, uncovering the asymmetric transfer across invariants and the location transfer effect within each invariant, both determined by structural stability. Specifically, learning effects of low-stability invariants transferred to those with higher stability, but not vice versa, with a similar pattern observed in DNNs. Additionally, location transfer effects were more pronounced for more stable invariants. These behavioral findings, alongside observed weight changes in DNNs, can be interpreted through a combination of the Klein’s Erlangen Program and RHT.

From the perspective of the Klein hierarchy of geometries, the asymmetric transfer can be expected given the nested relationship of the three geometrical invariants (***Klein, 1941***). On one hand, changes in a high-stability invariant include changes in less stable invariants, allowing improved discrimination abilities from training on low-stability invariants to assist in the discrimination of more stable ones. On the other hand, high-stability invariants are represented holistically during learning, making it difficult to encode their individual components in isolation. When performing a form discrimination task, the learners only need to extract the most stable invariants without necessarily extracting the embedded, less stable ones. This results in the suppression of less stable invariants when embedded within more stable configurations, limiting the generalization of learning from high-stability to low-stability invariants. Furthermore, the location generalization within each invariant and the distribution of learning across DNN layers suggest an underlying mechanism within the RHT framework. Specifically, training on high-stability invariants showed greater generalization to new locations (Experiment 3) and induced more weight changes higher in the DNN hierarchy (Experiment 4), indicating involvement of higher-level cortical areas with large receptive fields. And the order of weight changes across DNN layers during orientation task training aligns with the RHT prediction that higher visual areas change before lower ones (***Ahissar and Hochstein, 2004***; ***Hochstein and Ahissar, 2002***)

Taken together, we speculate that the VPL of high-stability invariants occurs earlier and relies more on higher-level cortical areas, while learning of low-stability invariants requires further optimalization and relies more on lower-level cortical areas with higher resolution for finer discriminations. Due to the feedforward anatomical hierarchy of the visual system (***Markov et al., 2013***), the modifications of lower areas caused by training on low-stability invariants will also affect higher-level visual areas, thus influencing the discrimination of high-stability invariants and resulting in transfer effects from low-to-high-stability invariants.

As stated in Introduction, there are several factors (difficulty, precision, variability, salience) which had been proven to determine the locus of plasticity and thus the generalization of learning. Specifically, low-difficulty, low-precision, high-salience, high-variability tasks are learned on the basis of neurons in the higher-order visual cortex, with involvement of more invariant neurons. Our demonstration that training on higher-stability invariants will lead to involvement of higher order visual areas is consistent with these findings in some degree, given that high-stability invariants possess both high variability (underwent more general shape-changing transformation according to definition) and high perceptual salience. However, it should be noted that difficulty and precision fail to explain the pattern of generalization within or across invariants observed in Experiment 2, 3 and 4 (as demonstrated in Results). Hence, the structural stability of form invariants needs to be considered as a more essential and determinant factor underlying the generalization of VPL, at least in our research.

Asymmetric transfer of VPL has also been reported in previous literature. For instance, Huang et al. found an asymmetric transfer of learning from motion discrimination to detection, explaining this phenomenon as a result of only task-relevant stimulus information being learned (***Huang et al., 2007***)—an explanation that aligns with our findings. According to the Klein hierarchy of geometries, the discrimination of high-stability invariant was a task-relevant prerequisite for discriminating low-stability invariant. For example, discriminations of collinearity and parallelism were implicitly required for the orientation task and thus benefited from orientation training. Conversely, during training on collinearity and parallelism, the specific orientations of the lines, while present but task-irrelevant, were not learned. A similar principle can be applied to explain the transfer from parallelism to collinearity. Additionally, previous evidence has demonstrated that precise discrimination learning could transfer asymmetrically to coarse discriminations (***Hung and Seitz, 2014***; ***Jeter et al., 2009***). However, precision could not explain the asymmetric transfer observed in our study, as elucidated in Experiment 2 and 4. Moreover, several studies have revealed the asymmetric transfer from clear to noisy display (***Chang et al., 2014***, ***2013***; ***Chen et al., 2016***; ***Dosher and Lu, 2005***), where VPL of a visual feature embedded in zero external noise, such as orientation, motion direction, or binocular disparity, transferred to the same feature in high external noise. Evidence from transcranial magnetic stimulation (TMS) studies in humans and physiological studies in monkeys suggests that this generalization results from functional reweighting, where training reweights the causal contributions of cortical areas involved in processing clear and noisy features (***Chang et al., 2014***; ***Chen et al., 2016***; ***Chowdhury and Deangelis, 2008***; ***Liu and Pack, 2017***). Specifically, training with clear stimuli containing informative local cues enhances reliance on lower-level cortical areas, where the optimized representation of these stimuli makes them more resistant to noise. This leads to a greater role of lower-level areas in perceptual decisions for noisy stimuli and supports the transfer of learning to such conditions. A similar reweighting mechanism may underlie the asymmetric transfer observed in our study, a possibility that future research should explore further.

According to the results from long-term training regimes in Experiment 2, invariants with lower-stability demonstrated greater learning effects and faster learning rates (***Figure 6—figure supplement 2***). The greater magnitude of learning and final layer change in DNNs (***Figure 9A, C***) for lower-stability invariants consistently suggest that less stable invariants exhibit greater plasticity. However, behavioral findings on learning rates contrast with DNN simulations, where performance of higher-stability invariants reached an asymptote earlier (***Figure 8B***). This inconsistency may be due to initial performance differences between human participants and DNNs. Previous research has shown that learning rate and magnitude of learning are negatively correlated with initial performance (***Yang et al., 2020***). In biological visual systems, more stable invariants are easier detected (***Chen, 2005***, ***1985***, ***1982***; ***Todd et al., 2014***, ***1998***) and demonstrate better initial performance (as demonstrated in the psychophysics experiments of this study), leading to lower learning rates and learning effects. In contrast, DNNs typically begin training with initial accuracies near chance level (50%) for all three invariants. We speculate that the learning process in DNNs may model not only VPL but also aspects of evolution or development, where more stable invariants with greater ecological significance mature earlier and thus undergo less improvement through VPL.

In conclusion, our findings on the learning and transfer effects of geometrical invariants align well with the Klein hierarchy of geometries. The results on location generalization and DNN simulations suggest that VPL of more stable invariants occurs earlier in higher-level visual areas, followed by VPL of less stable invariants in lower-level areas, consistent with the predictions of RHT. In light of Gibson’s theory of perceptual learning, we infer that the Klein hierarchy of geometrics belong to what Gibson referred to as “structure” and “invariant” (***Adolph and Kretch, 2015***; ***Gibson, 1970***). More importantly, invariants with higher structural stability parallel Gibson’s “higher-order invariants” (***Gibson, 1971***), which are extracted earlier in both perception and perceptual learning processes. Perceptual learning is a process of differentiation, beginning with the extraction of global, more stable invariants, and progressively involving local, less stable invariants to meet various task demands. Since the extraction of high-stability invariants is implicitly required in this process and can benefit from refined local information, the perceptual system which becomes more differentiated also develops an enhanced capacity for distinguishing high-stability invariants, leading to the asymmetric transfers observed in our research.

Future research should employ various neuroimaging and neurophysiological techniques to elucidate the neural mechanisms underlying VPL of geometrical invariants with varying stability and validate our theoretical framework. Such investigations could advance our understanding of object recognition, conscious perception and perceptual development.

## Methods and Materials

### Participants and apparatus

A total of 148 right-handed healthy subjects participated in this study: 44 in Experiment 1 (24 female, mean age 23.70 ± 3.46 years), 45 in Experiment 2 (26 female, mean age 23.02 ± 3.40 years), 21 in Experiment 2—long-term VPL (10 female, mean age 23.14 ± 1.85 years), 38 in Experiment 3 (20 female, mean age 24.39 ± 2.57 years). All subjects were naïve to the experiment with normal or corrected-to-normal vision. Subjects provided written informed consent, and were paid to compensate for their time. Sample size was determined based on power calculations following a pilot study showing significant learning effect of the collinearity task for effect size of Cohen’s d = 0.728 at 80% power. In each experiment, subjects were randomly assigned to one of three training groups (colli., para., and ori. training group). The study was approved by the ethics committee of the Institute of Biophysics at the Chinese Academy of Sciences, Beijing.

Stimuli were displayed on a 24-inch computer monitor (AOC VG248) with a resolution of 1920 × 1080 pixels and a refresh rate of 100 Hz. The experiment was programmed and run in MATLAB (The Mathwork corp, Orien, USA) with the Psychotoolbox-3 extensions (***Brainard, 1997***; ***Pelli, 1997***). Subjects were stabilized using a chin and head rest with visual distance of 70 cm in a dim ambient light room.

### Stimuli and tasks for psychophysics experiments

In Experiment 1, the stimulus arrays were designed to measure the learning effects of different geometrical invariants using the “configural superiority effects” (CSEs) paradigm (***Pomerantz et al., 1977***; ***Pomerantz and Portillo, 2011***). CSEs refer to the findings that global configural relations between simple components rather than the components themselves may play a basic role in visual processing and were originally revealed by an odd-quadrant discrimination task, as illustrated in ***Figure 2—figure supplement 1***. This paradigm was also adapted to measure the relative salience of different levels of invariants (***Chen, 2005***). ***Figure 2A*** represents discrimination tasks based on difference in different geometrical properties. ***Figure 2Aa*** represents a discrimination task based on a difference in collinearity, which is a kind of projective property. Here, three quadrants have lines with one bend along their length, whereas the odd quadrant has straight lines. ***Figure 2Ab*** represents a discrimination task based on a difference in parallelism, which is a type of affine property. The orientation of the line segments in the odd quadrant differed from that of the other quadrants. In ***Figure 2Ac***, the exact same component line segments were present in all quadrants; however, the odd quadrant contained arrows with different orientation, a type of Euclidean property. The stimuli are composed of white line segments with luminance of 38.83 cd/*m*^2^ presented on black background with luminance of 0.13 cd/*m*^2^. The stimulus array is consisted of four quadrants (visual angle 2.6° × 2.6° for each quadrant) and presented in the center region (visual angle, 6° × 6°). The subjects need to identify which quadrant is different from the others. Either of the two states of an invariant may serve as a target. For example, in the Euclidean invariant condition (***Figure 2Ac***), both upward and downward arrows could be the target. A green central fixation point (RGB [0,130,0], 0.15°) was presented throughout the entire block. Subjects were instructed to perform the actual task while maintaining central fixation. Each trial began immediately after the Space key was pressed. The stimulus array was presented until the subject indicated the position of the odd quadrant (“target”) via a manual button press as fast as possible on the premise of accuracy. The response time (RT) in each trial was calculated from the onset of the stimulus array. A negative feedback tone was given if the response was wrong.

The sample stimuli and the layout of a stimulus frame in Experiment 2 are illustrated in ***Figure 4*** and ***Figure 4—figure supplement 1***, respectively. Individual stimulus could occur in one of four quadrants, approximately 250 arc min of visual angle from fixation (*R*); two stimuli were presented at two diagonally opposite quadrants on each trial. The presentation diagonal was randomly selected with equal probability. The “target” referred to the pair of non-collinear lines (in other words, a line with one bend along its length) for the colli. task, the pair of unparallel lines for the para. task, and the more clockwise line for the ori. task. The trials not involved presentation of a “target” were set as catch trials (***Figure 4***). All stimuli are white (38.83 cd/*m*^2^) presented on black background (0.13 cd/*m*^2^). Each stimulus is composed of a group of line(s): a pair of lines which are collinear or non-collinear in colli. task, a pair of lines which are parallel or unparallel in para. task, and a single long line in ori. Task (***Figure 4—figure supplement 1***). The stimuli for the three tasks were made up of exactly the same line-segments. Thus, line-segments as well as all local features based on these line-segments, such as luminous flux, and spatial frequency components, were well controlled. The length of line (*l*) in colli. and para. task is 80 arc min, which is half of the length of line ori. task. The width of line is 2 arc min. The distance (*d*) between the pair of lines in the parallelism task is 40 arc min. The “base” orientation (the dashed line in ***Figure 4*** and ***Figure 4—figure supplement 1***) was randomly selected from 0-180°. For colli. and para., the “base” orientation for each stimulus on a stimulus frame was selected independently.

Schematic description of a trial in Experiment 2 is shown in ***Figure 5***, to make the first-order choice, subjects were instructed to press the J key if they thought there was a “target” as defined by each task, and they should press the F key if the contrary is the case (that is, there was not a bent line for the colli. task, there was not a pair of unparallel lines for the para. task, and the two lines were oriented in the same direction for the ori. task). A second-order choice needed to be made if subjects have pressed the J key in the first-order choice: they should select which one was the “target” and report its position by pressing the corresponding key (the F, J, V, and N keys correspond to the upper left, upper right, lower left, and lower right quadrants, respectively). The response of a trial was regarded as correct only if both choices were correct. This type of experimental design with two-stage choice options together with relatively higher proportion of catch trials (one third of all trials) wound help reduce false alarms and response bias.

The stimuli and tasks in Experiment 3 are identical to those in Experiment 2, with the sole difference being that the presentation diagonal was fixed during the same phase (either the training or testing phase).

### Procedure for psychophysics experiments

The overall procedure of Experiment 1 is show in ***Figure 2B***. The main VPL procedure consisted of three phases: pre-training test (Pre-test), discrimination training (Training), and post-training test (Post-test). During the testing phases, the three form invariant discrimination tasks were performed counterbalance across subjects at three blocks, respectively. During the training phase, all subjects were required to finish 10 blocks of the training task which is determined by their group. Each block contained 40 trials. Before all of the phases, subjects practiced 5 trials per task to make sure that they fully understood the tasks.

The procedure of Experiment 2 is similar to that of Experiment 1 except for no involvement of the baseline training. During each block, subject’s threshold was measured for each of the three tasks using a QUEST staircase of 40 trials augmented with 20 catch trials, leading to overall 60 trials per block. During the testing phases, the tests for the three tasks were counterbalanced across subjects. The training phase contained 8 blocks of the training task corresponding with the group of subjects. Each subject practiced one block per task with large angle difference to make sure that they fully understood the tasks.

Experiment 3 consisted of only two phases: discrimination training (8 or 9 blocks) and a transfer test (one block) on the same task, with stimuli presented at untrained retinal locations.

### Deep neural network simulations

The deep learning model used in this paper was adopted from Wenliang and Seitz (***Wenliang and Seitz, 2018***). The model was implemented in PyTorch (version 2.0.0) and consists of two parallel streams, each encompassing the first five convolutional layers of AlexNet (***Krizhevsky et al., 2017***) plus one fully connected layer which gives out a single scalar value. The network performed a two-interval two-alternative forced choice (2I-2AFC) task. One stream accepted one standard stimulus and the other stream accepted one comparison stimulus. The comparison stimulus was then compared to the standard stimulus. After the fully connected layers, the outputs of the two parallel streams – two scalar values – were entered to a sigmoid layer to give out one binary value indicating which stimulus was noncolinear, unparallel, or more clockwise, equivalent to the “target” in Experiment 2. Weights at each layer are shared between the two streams so that the representations of the two images are generated by the same parameters.

For the stimulus images, 12 equally spaced “base” orientations were chosen (0-165°, in steps of 15°), and the “base” orientation of the standard and comparison was chosen independently except for the orientation discrimination task. We trained the network on all combinations of the 3 parameters: (1) angle separation between standard and comparison —— ① 5° for colli. & para. and 10° for ori., ② 10° for colli. & para. and 20° for ori., ③ 20° for colli. & para. and 40° for ori.; (2) distance between the pair of lines for para. —— ① 30 pixels, ② 40 pixels; (3) the location of the gap on line for colli. —— ① the midpoint; ② the front one-third. So there were overall 12 (3 × 2 × 2) stimulus conditions. It should be noted that the angle separations in the orientation task were always twice the angle separations in the other two tasks in each condition, this proportion was based on the initial threshold observed in the behavioral experiment. Here are the other stimulus parameters: length of line in para. (100 pixels); length of line in colli. and ori. (200 pixels); width of line (3 pixels); radius of gap for colli. (5 pixels). Each stimulus was centered on an 8-bit 227 × 227 pixel image with black background.

Three networks were trained on the three tasks (colli., para., ori.) respectively, repeated in 12 stimulus conditions. Each network was trained for 1000 iterations of 60-image batches with batch size of 20 pairs, and meanwhile was tested on the other two untrained tasks. Learning and transfer performances were measured at 30 approximately logarithmically spaced iterations from 1 to 1000. We used the same feature maps and kernel size as the original paper. Network weights were initialized such that the last fully connected layer was initialized by zero and weights in the five convolutional layers were copied from an AlexNet trained on object recognition (downloaded from http://dl.caffe.berkeleyvision.org/bvlc_reference_caffenet.caffemodel) to mimic a (pretrained) adult brain. Training parameters were set as follow: learning rate = 0.0001, momentum = 0.9. The cross-entropy loss function was used as an objective function and optimized via stochastic gradient descent.

### Data analysis

All data analyses were carried out in Matlab (The Mathwork corp, Orien, USA) and Python (Python Software Foundation, v3.9.13). No data were excluded.

Human behavioral data were analyzed using analysis of variance (ANOVA), post-hoc tests, one-sample t-tests and paired t-tests. For ANOVAs and post-hoc tests, we computed 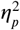 and Hedge’s g as effect sizes. For one-sample and paired t-tests, we computed Cohen’s d as effect size. To quantify learning effect, we computed the Learning Index (LI) (***Petrov et al., 2011***) as follows:

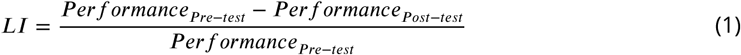

where *performance* refers to the RT or threshold obtained at the testing phases.

To quantify location transfer in Experiment 3, we computed the Specificity Index (SI) (***Ahissar and Hochstein, 1997***) as follows:

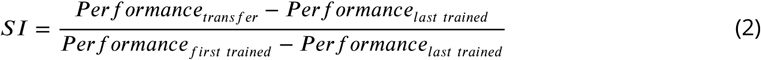

In DNN experiment, Ordinary Least Squares (OLS) method of linear regression was implemented to analyze the transfer effects between different tasks. The following equation describes the model’s specification:

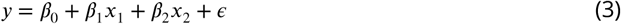

where *y* represents the dependent variable (final accuracy), *x*_1_ and *x*_2_ represent the two features (training and test task) respectively. *β*_0_ is the intercept, *β*_1_, *β*_2_ are coefficients of the linear model, and *∈* is the error term (unexplained by the model).

To estimate the learning effects in layers of DNN, the differences in weights before and after training at each layer were measured. Specifically, for a particular layer with *N* total connections to its lower layer, we denote the original *N*-dimensional weight vector trained on object classification as *w* (*N* and *w* are specified in AlexNet), the change in this vector after perceptual learning as *δw*, and define the layer change as follows:

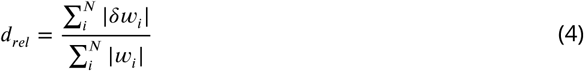

where *i* indexes each element in the weight vector. Under this measure, scaling the weight vector by a constant gives the same change regardless of dimensionality, reducing the effect of unequal weight dimensionalities on the magnitude of weight change. For the weights in the final read-out layer that were initialized with zeros, the denominator in Equation 1 was set to *N*, effectively measuring the average change per connection in this layer. Due to the convolutional nature of the layers 1–5, *d*_*rel*_ is equal to the change in filters that are shared across location in those layers. When comparing weight change across layers, we focus on the first five layers unless otherwise stated.

OLS method of linear regression with interaction terms was implemented to analyze the learning effects across layers. The following equation describes the model’s specification:

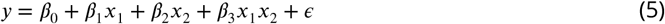

where *y* represents the dependent variable (PSI or final layer change), *x*_1_ and *x*_2_ represent the two features (training task and layer number) respectively. *β*_0_ is the intercept, *β*_1_, *β*_2_ and *β*_3_ are coefficients of the linear model, and *ε* is the error term (unexplained by the model).

## Code availability

The deep neural network is available from the original authors on GitHub (https://github.com/kevin-w-li/DNN_for_VPL).

## Acknowledgments

This work was supported by the Ministry of Science and Technology of China grant and the Young Elite Scientists Sponsorship Program by CAST.

## Additional information

## Author contributions

Yan Yang, Conceptualization, Data curation, Software, Formal analysis, Validation, Investigation, Visualization, Methodology, Writing – original draft, Project administration, Writing – review & editing; Zhentao Zuo, Tiangang Zhou, Conceptualization, Resources, Supervision, Funding acquisition, Methodology, Project administration, Writing – review & editing; Yan Zhuo, Conceptualization, Resources, Supervision, Funding acquisition; Lin Chen, Conceptualization, Resources, Funding acquisition, Methodology.

## Ethics

Human subjects: Informed consent, and consent to publish was obtained from each observer before testing. The study was approved by the ethics committee of the Institute of Biophysics at the Chinese Academy of Sciences, Beijin (reference number:2017-IRB-004).

---

**Figure 2—figure supplement 1.**
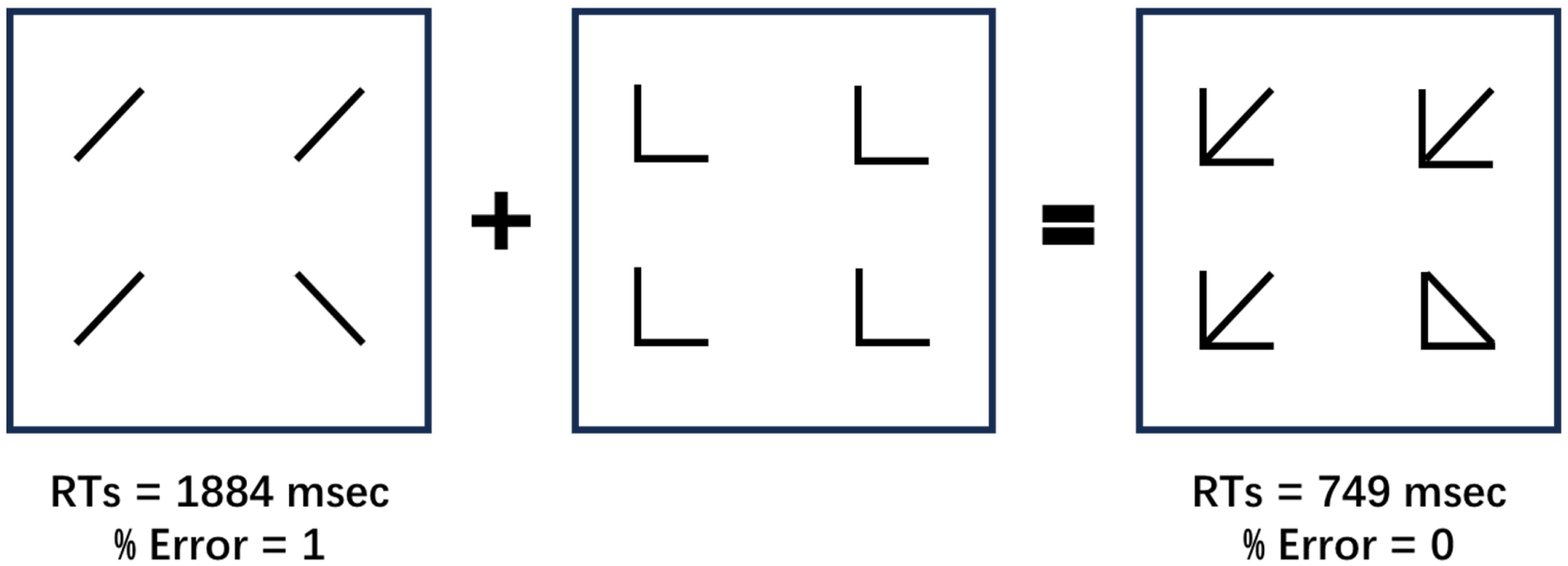
The configural superiority effect, adapted from (***Pomerantz et al., 1977***; ***Pomerantz and Portillo, 2011***). In the odd-quadrant discrimination task, observers were required to locate the odd stimulus in an array of four figures as quickly as possible. The left panel shows the *base display* where the odd quadrant differs from the rest in line slope. The center panel shows the *context display* with four identical quadrants. The *composite display* is shown in the right panel, which is simply the superposition of the base and context displays. Mean correct reaction times (RTs) and error rates are shown. Note that the reaction times and accuracies for the *composite display* were better than the *base display*.

**Figure 3—figure supplement 1.**
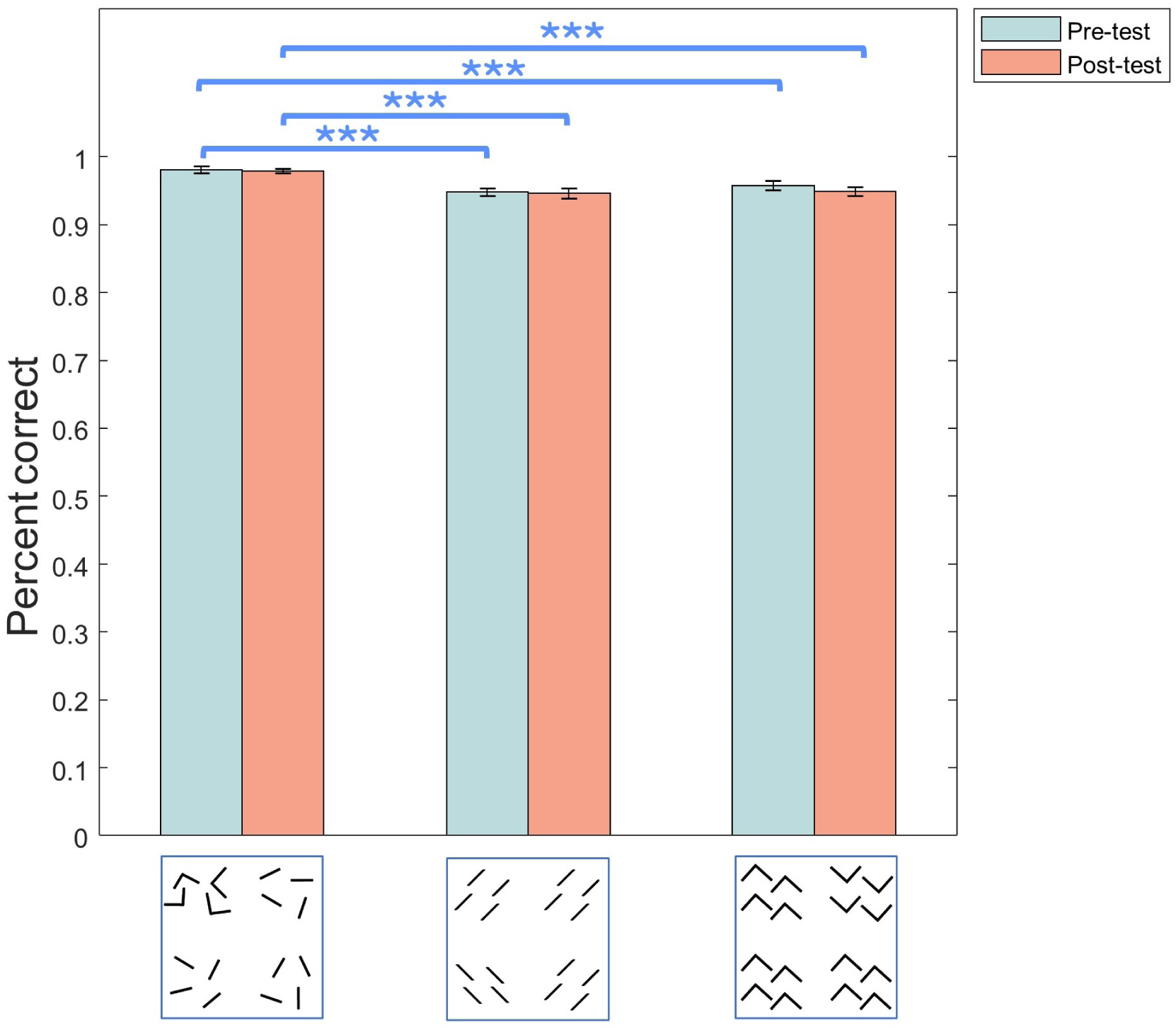
Accuracies for the three discrimination task measured at Pre-test and Post-test. Accuracy was defined as the average percentage correct per block. At Pre-test, the accuracies of the collinearity task were significantly higher than that in the parallelism (p < 0.0001) and orientation task (p = 0.0001). At Post-test, the accuracies of the collinearity task were still significantly higher than the other two tasks (p < 0.0001 for the parallelism task, p = 0.0001 for the orientation task). Statistical significance was calculated by paired t-test with FDR correction. (***p < 0.001, **p < 0.01, *p < 0.05). Error bars denote 1 SEM across subjects.

**Figure 3—figure supplement 2.**
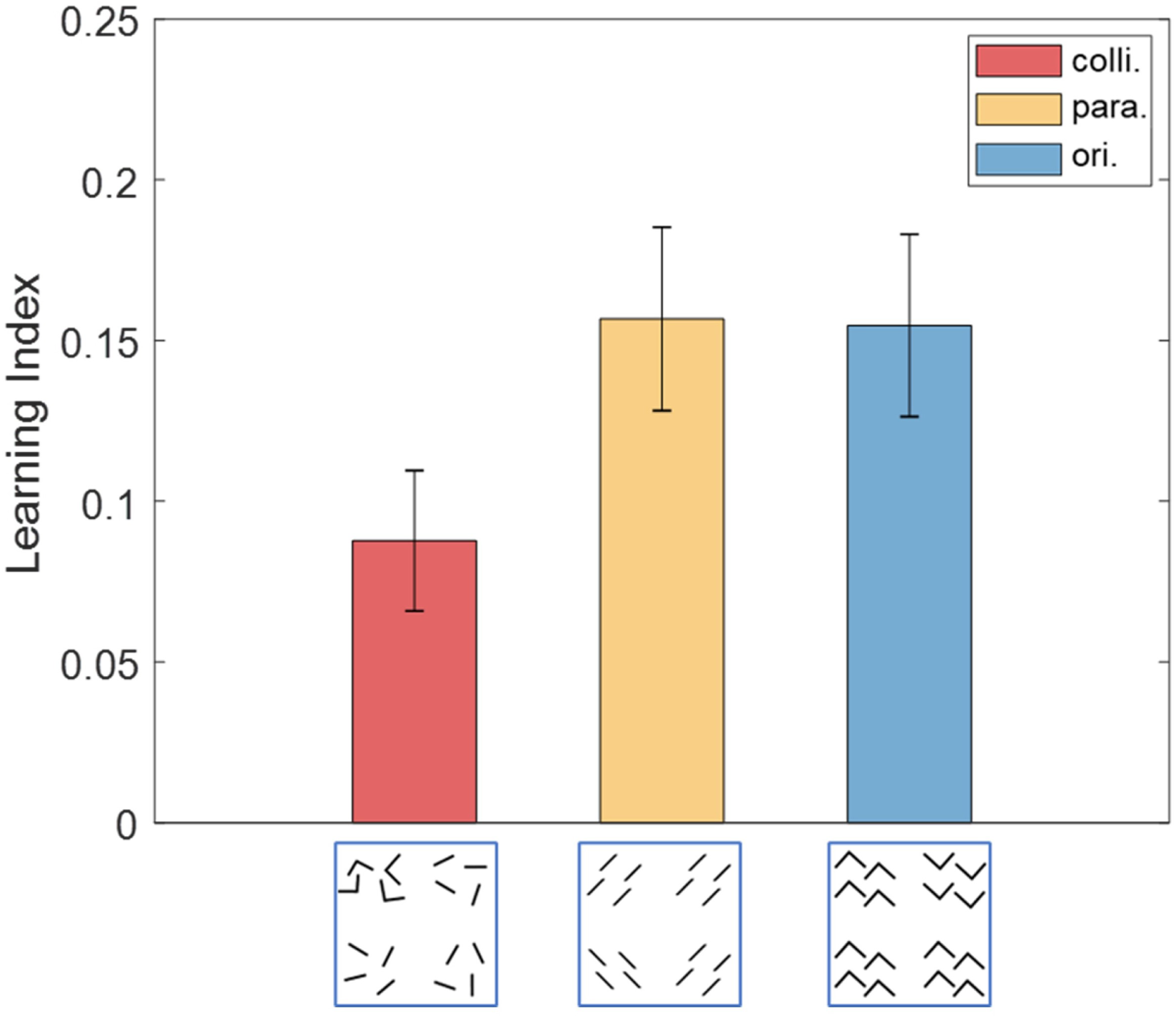
The learning indexes of the three geometrical invariants in Experiment 1. Error bars denote 1 SEM across subjects.

**Figure 4—figure supplement 1.**
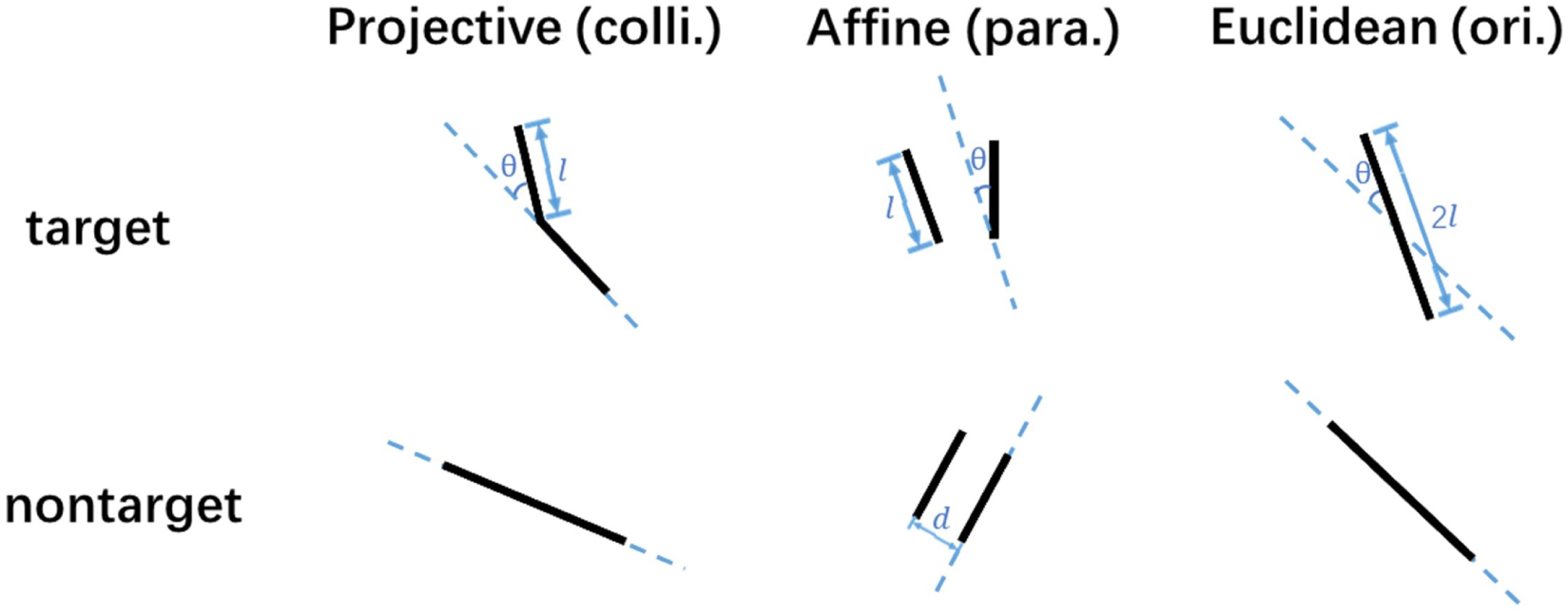
Examples of stimuli in Experiment 2. Sample stimuli in the collinearity (left), parallelism (middle) and orientation (right) discrimination task. The blue dashed lines represent the “base” orientation for each stimulus, and *θ* is the angle separation of the discrimination task.

**Figure 6—figure supplement 1.**
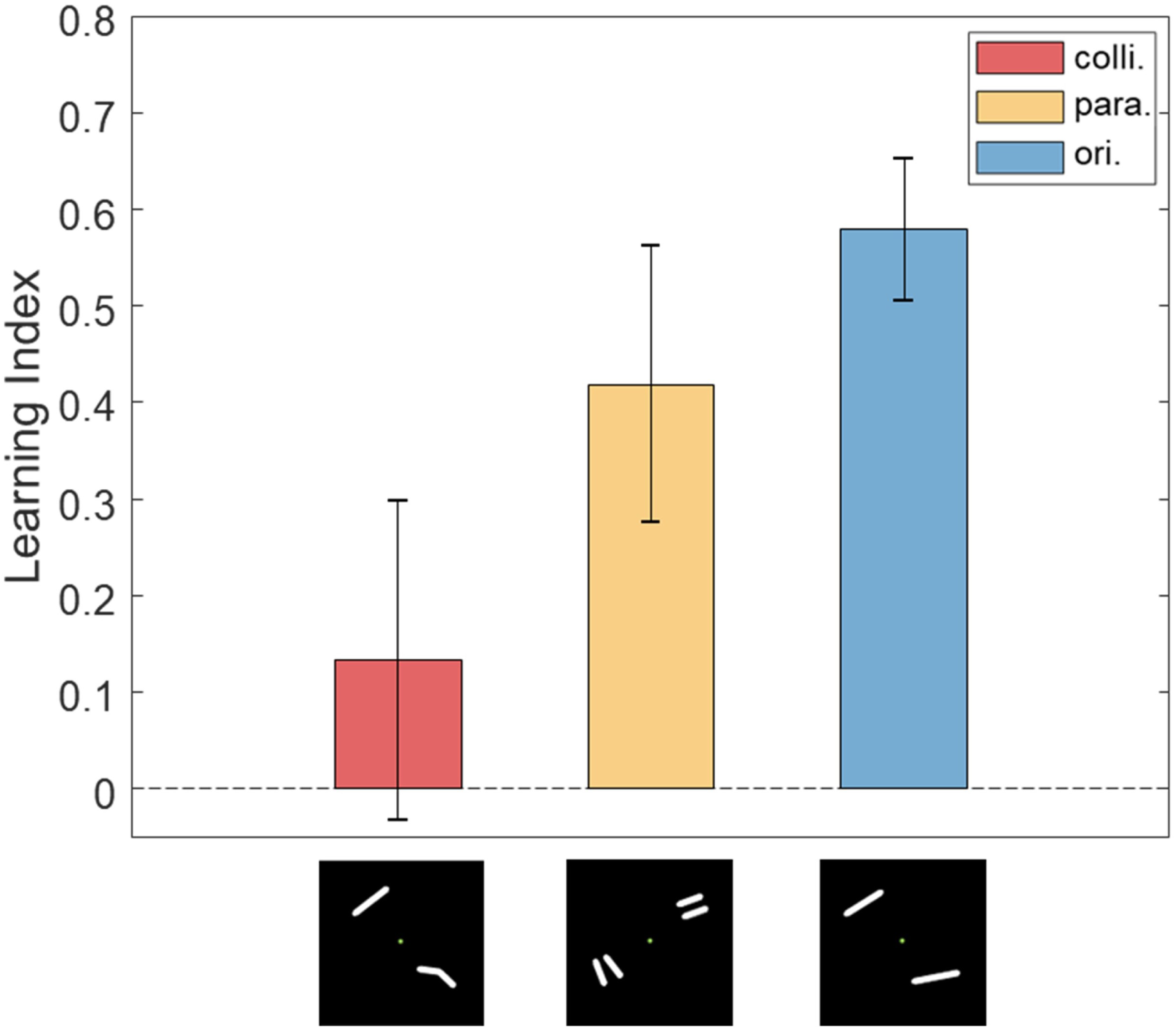
The learning indexes of the three geometrical invariants in Experiment 2. Error bars denote 1 SEM across subjects.

**Figure 6—figure supplement 2.**
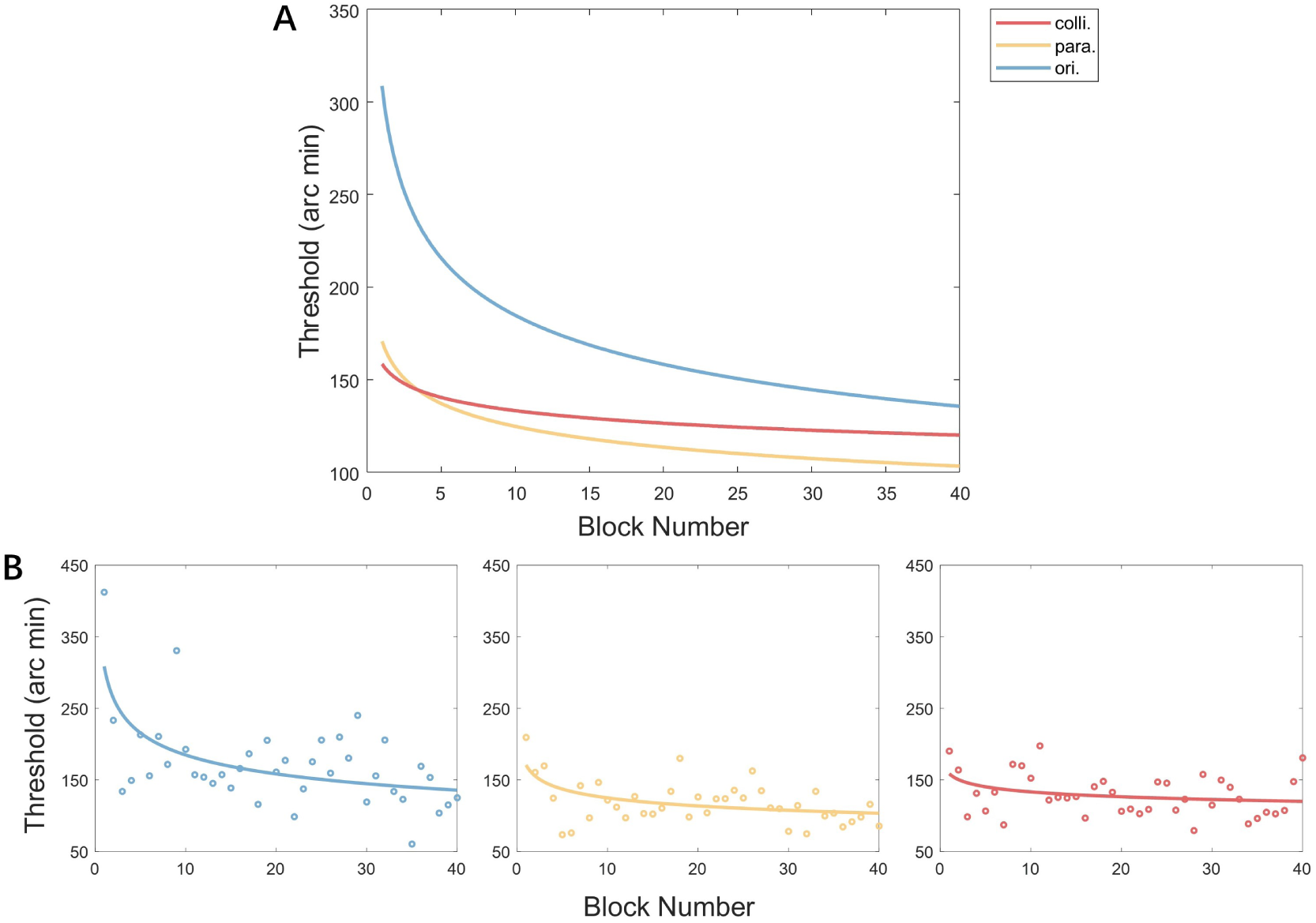
Bootstrap estimated learning curves across the three training tasks (n=7 for each task, the training lasted for 5 daily sessions, with each session consisting of 8 blocks). (A) Threshold data across blocks were fitted with a power function (*y* = *ax*^*b*^, where *y* is the block threshold) to estimate learning rate for each task. Subjects were randomly resampled with replacement 1,000 times, and parameters were fitted to each resample. Smooth curves were plotted based on the mean of the estimated parameters for each task. (B) Learning curves and performance across blocks for three training groups. Open circles indicate mean threshold per block.

**Figure 8—figure supplement 1.**
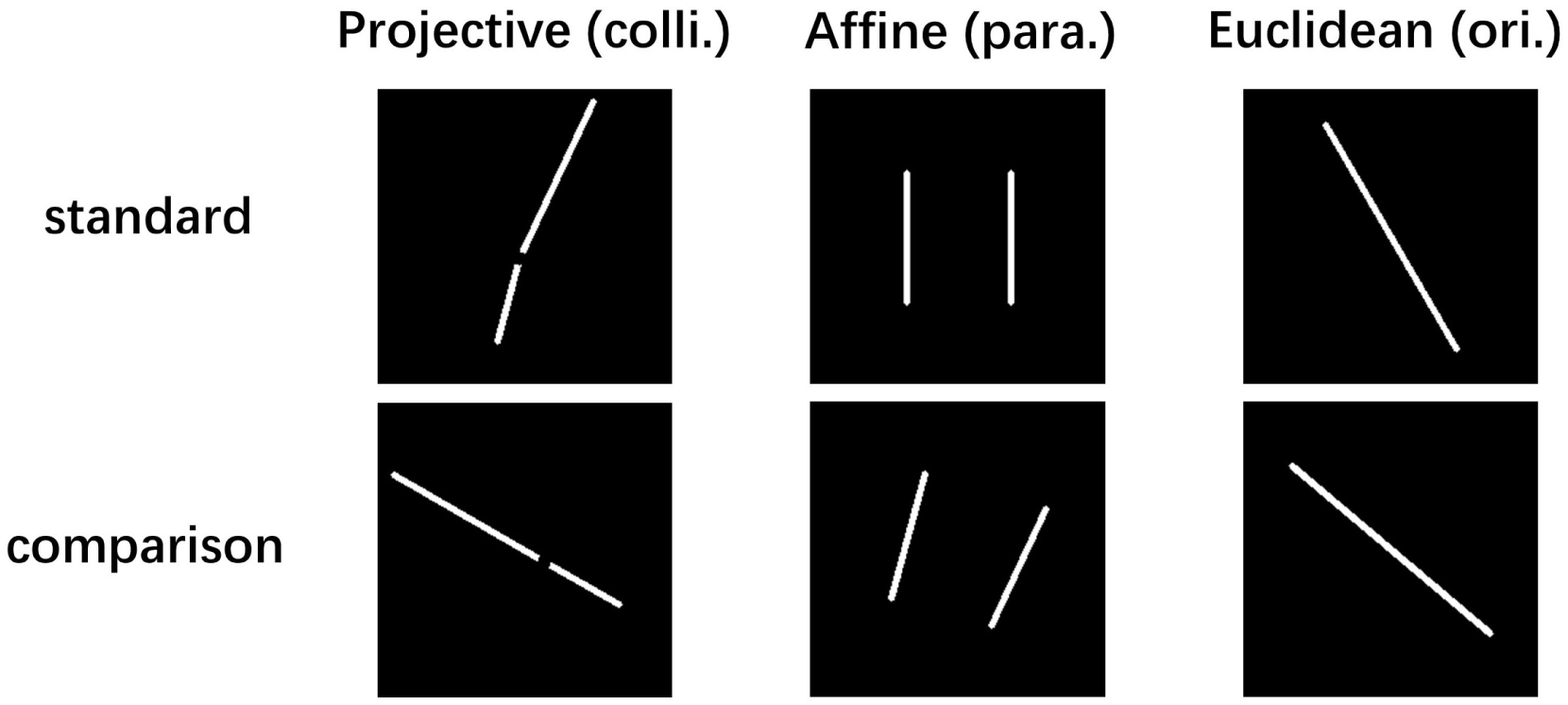
Stimulus examples in Experiment 3. Examples of the pairs of stimulus images for the three discrimination tasks in Experiment 3. The examples here are selected from the stimulus condition with the following parameters: angle separation (10° for colli. & para. and 20° for ori.), distance for para. (40 pixels), location of gap for colli. (the front one-third).

